# Cortical contributions to attentional orienting and response cancellation in action stopping

**DOI:** 10.1101/2024.11.08.622650

**Authors:** Sarah A Kemp, Sauro Salomoni, Pierre-Louis Bazin, Luke Pash, Rebecca J St George, Mark R Hinder

## Abstract

Action cancellation involves the termination of planned or initiated movement. Contemporary models of action cancellation, such as the Pause-then-Cancel model, propose that this occurs via a two-stage process, initiated in the cortex by the pre-supplementary motor area (preSMA) and inferior frontal gyrus (IFG). Previous experimental work using electromyography (EMG) has identified that the cancellation of actions can involve the partial activation of the responding muscles, which does not result in an overt behavioural response. In this study, we used functional near-infrared spectroscropy (fNIRS) to investigate the neural correlates of these partial responses in a modified stopping task (a response- and stimulus-selective stop-signal task), controlling for the attentional effects that have long confounded action cancellation research by comparing responses to stop stimuli with those to ignore stimuli. We identified stopping-related activity in the preSMA but not the IFG, consistent with predictions of the Pause-then-Cancel model. Additionally, we observed increased preSMA activity in trials where no partial responses occurred, potentially due to the cumulative effect of different inhibitory processes in those trials. This study also highlights the utility of combining fNIRS and EMG in examining the cortical correlates and dynamic processes involved in action cancellation.

## 1. Introduction

Movement adaptation is underpinned by response inhibition mechanisms, which coordinate in order to realise adaptive, goal-directed movement (Logan and Cowan, 1984). An important part of response inhibition is action cancellation, the sudden termination of planned or already-initiated movement in response to unexpected stimuli (Logan, 1981; Sebastian et al., 2018). As well as being a crucial part of everyday motor control, action cancellation is impaired in a number of neurological and psychiatric conditions, including Parkinson’s disease, attention-deficit hyperactivity disorder, obsessive-compulsive disorder, post-traumatic stress disorder, and schizophrenia (Benis et al., 2016; Borst et al., 2024; Hughes et al., 2012; Kibleur et al., 2016; Senkowski et al., 2023), and declines in older age (Healey et al., 2024; Kemp et al., 2024). Action cancellation is thought to be mediated by two cortico-basal ganglia pathways (the hyperdirect and indirect pathways; Jahfari et al., 2011; Schroll and Hamker, 2013), but the specific roles of these pathways and the neural mechanisms underlying action cancellation remain subjects of ongoing debate.

The indirect and hyperdirect pathways originate in the cortex and involve various basal ganglia regions, ultimately sending commands to effector muscles via the primary motor cortex (M1; Bingham et al., 2023; Schroll and Hamker, 2013). The indirect pathway is thought to originate in the pre-supplementary motor area (preSMA) and involve a broad range of basal ganglia areas, while the hyperdirect pathway is thought to originate in the inferior frontal gyrus (IFG), and include only the subthalamic nucleus (STN) and the globus pallidus interna (Bingham et al., 2023; Coudé et al., 2018; Nambu et al., 2002). Neuroimaging and brain stimulation studies have consistently implicated both the preSMA and IFG in action cancellation (see e.g., Boen et al., 2022; Borgomaneri et al., 2020; Forstmann et al., 2008; Gavazzi et al., 2019; Lee et al., 2016), but there is little consensus on their individual contributions to this process.

Action cancellation is often investigated using an SST, where participants make a standard response in most trials (e.g., a button press) following a ‘go’ signal. On a subset of trials (generally 25-33%), a stop signal will appear shortly after the go signal, requiring that the participant try to cancel their response (for a consensus paper regarding the use of the SST, see Verbruggen et al., 2019). However, the infrequent stop stimulus intrinsic to the SST is *unexpected* as well as requiring action cancellation. Thus, interpretation of experimental data and associated brain imaging using this paradigm may be confounded by attentional orienting, with stop trials engaging both action cancellation and attentional processes (Hampshire, 2015; Sharp et al., 2010). To control for these effects, some experimental work has used stimulus-selective SSTs, with additional infrequent stimuli that do not require action cancellation (e.g., Boehler et al., 2010; Chatham et al., 2012; Weber et al., 2023). In such paradigms, the preSMA has consistently been found to be more uniquely associated with action cancellation, with increased activity in successful stop trials (where action cancellation is required) compared to ignore trials (which involve attentional orienting but not action cancellation; Li et al., 2006; Rae et al., 2014; Sharp et al., 2010). Conversely, IFG activation is evident in both ‘stop’ *and* ‘ignore’ contexts, suggesting that its role in action cancellation may be more related to attentional orienting mechanisms than action cancellation *per se* (Boehler et al., 2010; Sharp et al., 2010).

Converging lines of evidence suggest that action cancellation occurs in two stages, underpinned by the hyperdirect and indirect pathways. Based on evidence from the rodent basal ganglia (Frank, 2006; Schmidt and Berke, 2017), these findings have recently been adapted into human action stopping in the Pause-then-Cancel (PTC) model (Diesburg and Wessel, 2021). The PTC model integrates evidence from a range of modalities and is rapidly emerging as one of the dominant models in action cancellation (Hannah et al., 2022; Pani et al., 2022; Tatz et al., 2021). Under this framework, upon presentation of an unexpected stimulus, the IFG sends a rapid ‘pause’ signal to M1 via the hyperdirect pathway. This dynamically increases the response threshold, putatively delaying the execution of movement without modifying it (Diesburg and Wessel, 2021). This occurs in response to infrequent or unexpected stimuli, regardless of an imperative to stop. The manner in which the ‘pause’ mechanism alters response threshold is proposed to be generic and non-selective, modifying excitability in effector muscles both relevant and irrelevant for the movement (Diesburg and Wessel, 2021; Wadsley et al., 2023). The pause is followed by a ‘cancel’ process, which modifies the movement according to contextual requirements once the stimulus has been fully interpreted. This slower mechanism is proposed to occur via the indirect pathway (initiated in the preSMA) and to be specific to action stopping or action modification (Tatz et al., 2021). However, the majority of this evidence comes from contexts involving global stopping paradigms (see below); the extent to which these processes apply in other stopping contexts and how the neural correlates may differ across paradigms is yet to be determined.

The majority of SSTs use non-selective stopping paradigms, where the participant makes a choice response in each trial (e.g., a left or right unimanual button press) and attempts to cancel the entire movement in a stop trial, resulting in no behavioural response (Verbruggen et al., 2019). Whilst these paradigms are informative, real-world stopping scenarios are often more complex, involving the termination of only specific components of an initiated movement. This can be assessed using response-selective SSTs, where participants cancel only a *part* of the initiated action (e.g., a bimanual button press is required following the go signal, with movement from one hand only stopped in response to the stop signal; Coxon et al., 2007; Salomoni et al., 2023; Weber et al., 2023). It has been consistently observed that following this selective stopping, the continuing behavioural response is markedly slower than reaction time in standard go trials (referred to as the stopping delay or stopping interference effect; for a review, see Wadsley et al., 2022), but there is, as yet, no consensus regarding how these more complex stopping scenarios are handled at a neural level (Aron and Verbruggen, 2008; Coxon et al., 2007; Hannah et al., 2022; Raud et al., 2020).

Electromyography (EMG) is ideal for capturing the electrophysiological changes that occur during movement execution and action cancellation (Raez et al., 2006). Importantly, it can be used to detect partial responses: instances where EMG amplitude increases above a certain threshold without resulting in an overt behavioural response (also referred to as prEMG or partial bursts; Raud et al., 2022; Salomoni et al., 2023; Thunberg et al., 2024). Observed in some successful stop trials, these partial responses are thought to reflect inhibition at the muscle, where an initiated action is terminated before producing a behavioural response (Atsma et al., 2018; Raud et al., 2022). Partial responses thus represent dynamic changes that can be measured with high temporal sensitivity, allowing important distinctions to be made between trials with no behavioural output (MacDonald et al., 2017; Wadsley and Greenhouse, 2024). The precise neural underpinnings of these dynamic changes are unknown and particularly, an open question remains regarding whether neurological distinctions can be made between trials with and without partial responses.

Functional near-infrared spectroscropy (fNIRS) is a non-invasive, optical neuroimaging technique that estimates concentration changes in oxygenated (Hb) and deoxygenated haemoglobin (dHb; Pinti et al., 2020). Hb and dHb differentially absorb near-infrared light, meaning fluctuations in their concentration alters the fNIRS signal (Rahman et al., 2020). Changes to Hb and dHb concentration are associated with neural activity, a relationship which forms the basis for the blood-oxygenation level-dependent (BOLD) signal fundamental to functional magnetic resonance imaging (fMRI; Heeger and Ress, 2002). fNIRS is a more portable and convenient neuroimaging technique than fMRI, but has limited depth sensitivity, meaning it is ill-suited for examining subcortical processes (Pinti et al., 2020). However, it is ideal for examining cortical activation patterns and brain regions relatively close to the skull, such as the preSMA and IFG.

Here, we investigated some of the neural and electrophysiological processes associated with action cancellation using a response-and stimulus-selective SST. We measured cortical activity using fNIRS and electrophysiological activity in the effector muscles using EMG. Participants responded to go signals with bimanual button presses. When presented with a stop signal, they attempted to cancel the response of one hand while continuing with the other. Infrequent ignore signals (which bore no imperative to stop) were used to delineate the behavioural and neurophysiological changes associated with stopping versus attentional capture. Partial responses enabled us to identify trials in which the original bimanual response had been initiated but interrupted following the presence of a stop or ignore signal.

## 2. Methods

### 2.1. Participants

Thirty participants (mean age = 28.33 years, *SD* = 7.33, all right-handed, 19 females) were recruited via the University of Tasmania’s psychology research participation system or personal invitation. All participants had normal or corrected-to-normal vision, no colourblindess, and no history of neurological or major psychiatric disorders (including ADHD, due to alterations in inhibitory control; Senkowski et al., 2023). As compensation for their time, participants received either two hours of research credit or a $20 AUD shopping voucher. All participants gave written informed consent. One participant exhibited a proportion of successful stop trials (78.67%) that was over the recommended range (Verbruggen et al., 2019) and was thus excluded from the analysis. The fNIRS data of another participant was not usable due to a technical error. The final sample size was thus 29 for the behavioural and EMG components of the analysis, and 28 for the fNIRS component. The study was approved by the Tasmanian Human Research Ethics Committee (#H0014865) and conducted in accordance with the Declaration of Helsinki.

### 2.2. Stop-signal task (SST)

A response-and stimulus-selective SST was used, with go trials, ignore trials, and stop trials. Each trial began with a fixation cross, the duration of which was randomly drawn from a truncated exponential distribution (range: 500-2500 ms, median duration ≈ 1130 ms). This randomisation reduced the predictability of the go signal onset, ensuring participants were responding to the go signals themselves and not responding prematurely in anticipation (Verbruggen et al., 2019). The go signal was two green arrows. When they appeared, participants would make a bimanual, inwards button press (simultaneously with the left and right index finger) against vertically-mounted buttons as quickly as possible (see Figure 1). The response window was 2000 ms. In a stop trial, one of the green arrows would turn orange at a precise delay time after the go onset, indicating that the participant should withhold their response with that hand but still execute a button press with the other hand (response-selective inhibition). In an ignore trial, one of the arrows would turn purple after a delay. Here, participants were required to ignore the colour change, and make the standard bimanual response (stimulus-selective inhibition). Feedback on trial success and response time was then presented for 2000 ms.

**Figure 1.**
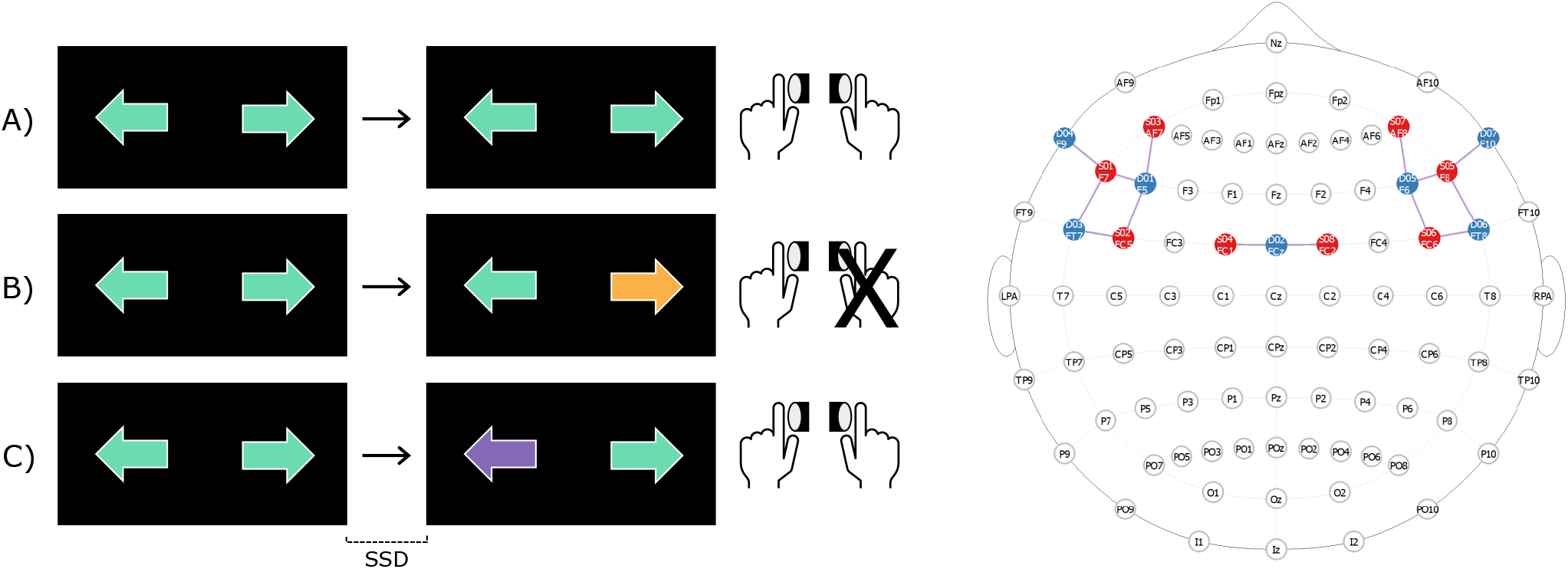
Schematic of the experimental task (left) and the custom montage used in the fNIRS set up (right). *Task*: Trials could be either go trials, stop trials, or ignore trials. Each trial started with the presentation of two green arrows, to which participants made fast-as-possible bimanual responses. In a go trial (A), the arrows remained green throughout the trial. In a stop trial (B), one of the arrows turned orange, indicating participants should try to cancel that component of the response and give a unimanual response. In an ignore trial (C), one of the arrows turned purple. Participants were to ignore this colour change and continue the bimanual response. The stop-signal delay (SSD) determined the onset of the colour change in each stop and ignore trial. Note that either arrow could change colour in a given trial, but only a right-sided stop trial (B) and a left-sided ignore trial (C) are illustrated here. *fNIRS*: Custom montage used to capture changes in haemoglobin concentration in the left and right inferior frontal gyri (LIFG and RIFG) and pre-supplementary motor area (preSMA). The montage comprised 14 channels (8 sources, 7 detector). Six channels were used for the LIFG, six for the RIFG, and two for the preSMA.

The latency between the onset of the go and stop signals (stop signal delay; SSD) was modified after each trial using a staircase procedure (Verbruggen et al., 2019). If a participant stopped successfully in a stop trial (i.e., gave the correct unimanual response), the SSD was increased by 50 ms on the next trial. Conversely, if a participant failed to stop the bimanual response or gave the wrong response (i.e., stopped the wrong hand), the SSD was *decreased* by 50 ms, making it easier for the participant to inhibit their response in the next stop trial. This iterative adaptation ensured that each participant was able to appropriately inhibit their response on approximately 50% of trials (Verbruggen et al., 2019). SSDs were tracked independently for each hand. The onset of the ignore signal was determined by the current SSD for that hand.

### 2.3. Procedure

Participants were seated approximately 80 cm from a computer monitor, with their forearms pronated and arms resting on a desk, shoulder width apart. Each index finger rested against a custom-made response button. The buttons were mounted in the vertical plane, such that pressing a button required participants to abduct their index fingers (Figure 1). This maximised activation of the first dorsal interrosei (FDI) muscles, as they act as agonists in this action (this also improves the signal-to-noise ratio [SNR] and the detection of EMG bursts; see 2.3.1). Button presses were registered via the Black Box Toolkit USB response pad and recorded using PsychoPy3 and Signal4 (Cambridge Electronic Design Ltd; Peirce et al., 2019). The SST was run via custom written code in PsychoPy3 (Peirce et al., 2019).

After receiving instructions, participants completed 20 practice trials, and then 680 trials divided in ten blocks (68 per block). Of these 680 trials, 450 were go trials (66%), 150 were stop trials (22%; 75 left and 75 right), and 80 were ignore trials (12%; 40 left, 40 right). Each 68-trial block contained the same proportions of trials, i.e., 45 go trials, 15 stop trials, and 8 ignore trials. After each block, participants had an opportunity to rest before continuing. The first six participants completed more trials than the rest of the cohort; it was found that the overall length of the experiment setup was longer than had been estimated during piloting, and trial numbers were thus reduced to minimise any possible fatigue effects. After careful inspection of the dataset, it was determined that these six participants did not differ from the remainder of the cohort in terms of their performance on stop and ignore trials, and had a similar reaction time (RT) distribution, indicating they performed similarly despite the longer experiment. All data from these participants were therefore retained in the analysis to maintain statistical power.

In addition to feedback at the end of each trial, participants received feedback at the end of each block, telling them their mean RT for that block. If the mean RT was more than 100 ms slower than in the previous block, additional feedback was provided, reminding them to respond as quickly as possible. This was to minimise proactive slowing strategies, as these create problems in the analysis of stop-signal data (Verbruggen et al., 2019) and are associated with distinct neural correlates compared to fast-as-possible responses (Sánchez-Carmona et al., 2016), especially in inferior frontal regions (Messel et al., 2019).

#### 2.3.1. Electromyography (EMG)

EMG was recorded using disposable adhesive electrodes (Ag/AgCl). Two electrodes were positioned in a belly-tendon montage over the FDI of each hand, with a ground electrode placed on the head of the ulna. Recordings were synchronised via a TTL pulse from the PsychoPy software, and made using the software Signal4 (Cambridge Electronic Design Ltd). The analogue EMG signals were bandpass filtered at 20-1000 Hz, amplified 1000 times, sampled at 2000 Hz (CED Power 1401 and CED 1902, Cambridge, UK), and saved into a PC for offline analysis.

#### 2.3.2. Functional near-infrared spectroscopy (fNIRS)

Near-infrared light intensity changes in the left IFG (LIFG), right IFG (RIFG), and preSMA were recorded with a custom fNIRS montage. Recordings were made with NirStar v15.3 with a sampling rate of approximately 7.82 Hz. Synchronisation was achieved using a digital TTL signal from PsychoPy. The system (NirSport, NIRs Medizintechnik GmbH, Berlin) emits near-infrared light at 760 − 850 nm wavelengths to capture haemodynamic changes in the cortical tissue up to 1.5 cm deep (Brigadoi and Cooper, 2015; Pinti et al., 2020).

The locations of the regions of interest (ROIs) were determined using masks from the Automated Anatomical Labeling 2 (AAL2) atlas. The AAL2 provides parcellation of the orbitofrontal cortex based on the Montreal Neurological Institute (MNI) standardised and spatially-normalised brain atlas (Rolls et al., 2015; Tzourio-Mazoyer et al., 2002). The montage was determined using the fNIRS Optodes’ Location Decider (fOLD) toolbox v2.2 (Zimeo Morais et al., 2018) and comprised 14 channels, with eight sources and seven detectors (see Figure 1). Six channels were used to capture the LIFG, six for the RIFG, and two for the preSMA. Given the small number of channels used to capture the preSMA, the limited spatial resolution of fNIRS, and the proximity of the preSMA to the midline (Hiroshima et al., 2014), this region was not split into right and left hemisphere. The channels and their approximate coordinates in MNI space (estimated using fOLD) are given in Table 1.

**Table 1.**
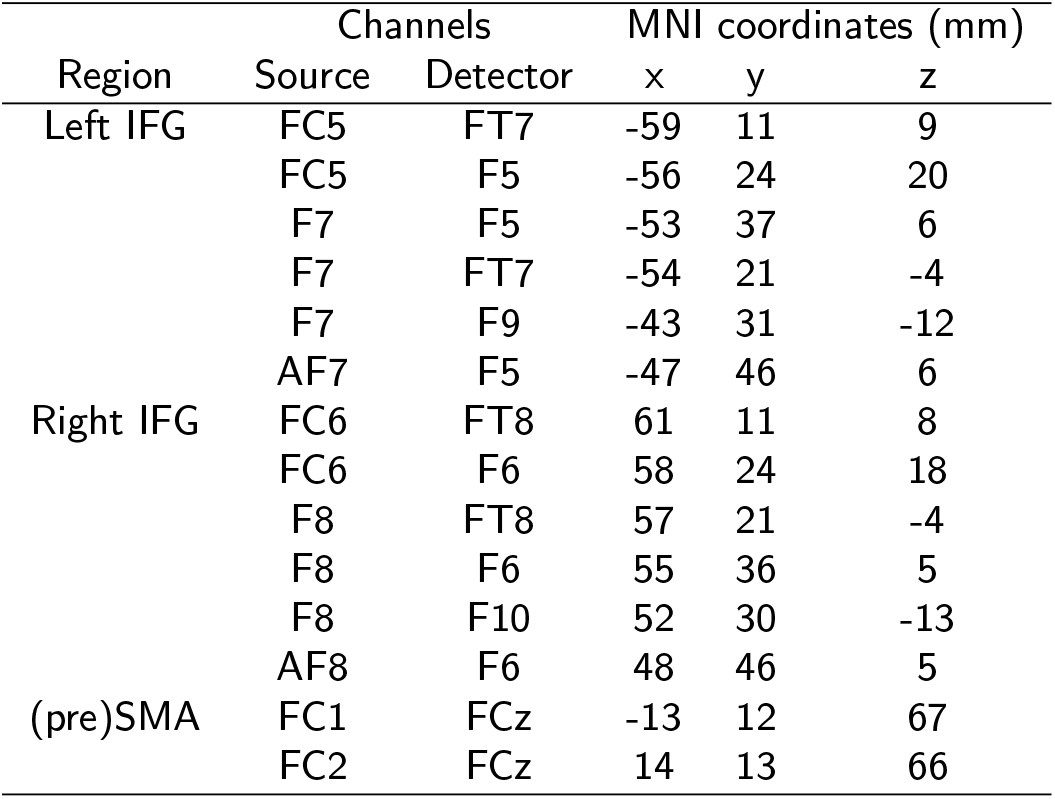
The custom montage given with the appoximate MNI coordinates for each channel. Montage was determined using the fNIRS Optodes’ Location Decider (fOLD) toolbox (Zimeo Morais et al., 2018).

### 2.4. Data Preprocessing

#### 2.4.1. EMG data

EMG preprocessing was performed using custom scripts, implemented in MATLAB R2017B (MathWorks, 2017). The signals were filtered using a fourth-order bandpass Butterworth filter at 20-500 Hz. To detect the precise onset and offset times of the EMG responses, we used a single-threshold algorithm as per Hodges and Bui (1996). A sliding window of 500 ms was used to find the segment in the timeseries with the lowest root mean squared amplitude, which was then used as a baseline to estimate the latencies of EMG responses. Signals from each trial were rectified and smoothed by lowpass filtering at 50 Hz to obtain EMG envelopes, which were used to esimate the amplitude of each burst. EMG points of interest are depicted in Figure 2.

**Figure 2.**
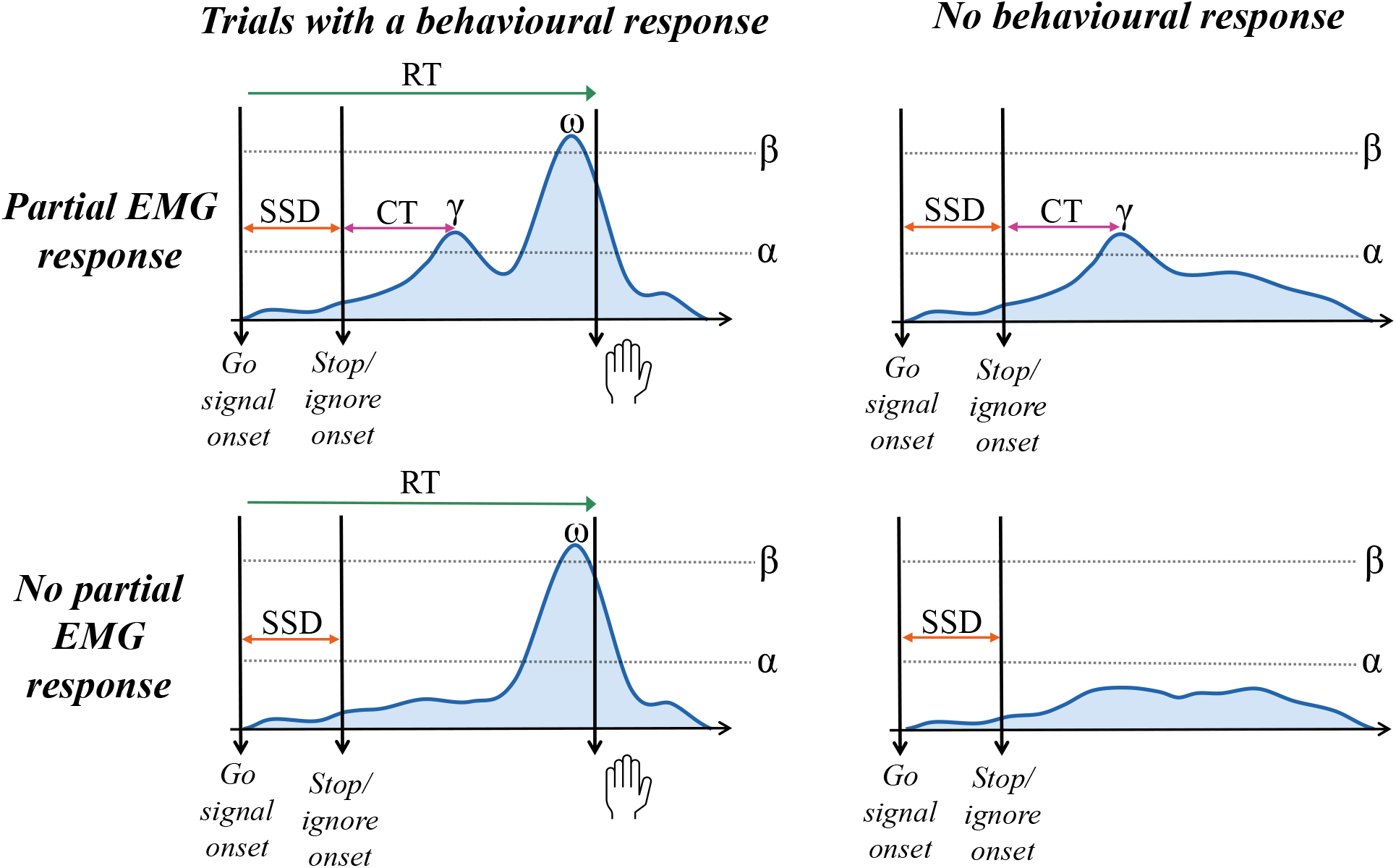
An illustrative schematic of EMG traces after presentation of a stop or ignore signal, showing EMG measures of interest. The onset of the stop or ignore signal is determined by the stop-signal delay (SSD). The reaction time (RT) is the time between onset of the go signal and the button press. Partial responses are instances where the EMG signal crosses the threshold *α* without having a subsequent button press. Response-generating EMG bursts are the increase in EMG closest to the recorded button press that cross the threshold *β*. Measures of interest include cancel time (CT; the latency between stop/ignore signal onset and the peak of the partial response *γ*), RT, and the amplitude of the response-generating EMG burst (*ω*).

EMG responses were categorised as instances where the amplitude of the smoothed signal exceeded a threshold of standard deviations (SD) above baseline (α in Figure 2). The responses were considered either *response-generating*, where this signal increase preceded an overt button press, or *partial responses*, where the signal crossed the threshold but did not result in a button press. Onset and offset times were also considered in the identification of these responses. Response-generating EMG bursts were defined as the last burst prior to the recorded button press (indicated by ω in Figure 2). Similar to our previous studies (Salomoni et al., 2023; Weber et al., 2023), partial EMG responses were identified as the first instance of the signal crossing the threshold in a trial (γ in Figure 2), as long as this increase occurred after the SSD and the amplitude of this response was greater than 10% of the average peak of the successful go trials for that participant. Partial response onset had to occur before any subsequent response-generating bursts in that trial; any partial movements close to the button press (simultaneous with a response-generating burst in the responding hand) are likely attributable to mirror activity or, if occurring after the press, due to other activation unrelated to the task. EMG responses separated by less than 20 ms were merged together, as these are unlikely to represent distinct activations. For each response detected (response-generating and partial responses), we extracted the times of the onset, peak, and offset. The latency of the peaks for partial responses and response-generating bursts, as well as the amplitude of the response-generating bursts, were used as measures of interest in the analysis (see Figure 2 and 2.5.2).

#### 2.4.2. fNIRS data

fNIRS data were processed using Homer3 v1.80.2, implemented in MATLAB R2017b (Huppert et al., 2009; MathWorks, 2017). In the processing stream, missing values were replaced by spline interpolation of the existing values using the PreprocessIntensity_NAN Homer3 function. PruneChannels was then used to remove channels with an SNR of below 2. Of the 14 channels in the montage, six participants had one noisy channel (resulting in the removal of 7% of their data), two participants had two noisy channels (14% of their data removed), and one had three (21% removed). The data were then converted from light intensity to optical density using the Intensity2OD Homer function, and motion artefacts removed using MotionCorrectPCArecurse. High-and low-pass filtering (0.005 Hz and 0.5 Hz, respectively) was applied using a third-order bandpass Butterworth filter via BandpassFilt. The optical density data were then converted to concentrations of Hb and dHb by the modified Beer-Lambert law using OD2Conc. A configuration file showing all functions and values for the processing stream can be found at https://osf.io/ztb5m/.

Increases in neural activity creates a subsequent increase in the haemodynamic response, influencing both Hb and dHb concentration (Heeger and Ress, 2002; Watanabe et al., 2017). Data analyses focused on Hb concentration, as this measure has a higher SNR than dHb and is most correlated with the fMRI BOLD response (Eggebrecht et al., 2012; St George et al., 2021, 2022). For each participant, the channels corresponding to each region (preSMA, RIFG, LIFG) were averaged to obtain a single timeseries per ROI. Trial-level timeseries were then extracted over a 10-second time window for each trial, starting 500 ms before go onset, resulting in a waveform of haemodynamic (Hb concentration) change for each trial per ROI. Figure 3 shows an example of the waveforms of a single participant, with Hb concentration changes in the three ROIs for a sample of go, failed stop, and successful stop trials.

**Figure 3.**
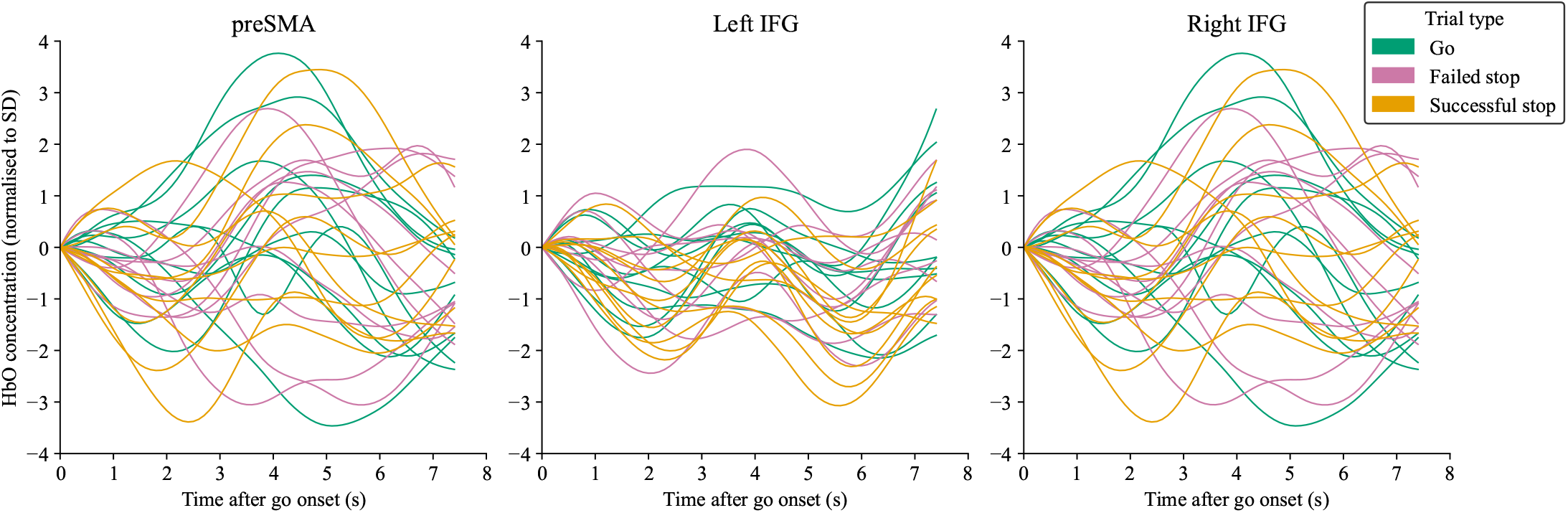
Sample trials from a representative participant, showing changes in Hb concentration in the preSMA and left and right IFG in the 8 seconds following go signal onset (time 0). Changes in neural activity triggered by the stimulus are reflected in Hb concentration changes, typically appearing with a 3-6 second delay. For analysis, the peak Hb concentration (local maximum) in the 2-7 seconds following go signal presentation was extracted per trial.

To assess differences in neural activity between different trial types, standard fNIRS approaches generally involve averaging the waveforms across trials. Values can then be extracted for each condition, such as the peak Hb concentration, mean Hb concentration, or increase in Hb concentration over a particular time period (see e.g., Healey et al., 2024; Kim et al., 2024; St George et al., 2021). Here, rather than averaging across trials, we extracted a value of Hb concentration in *each* trial. This enabled us to apply hierarchical (multi-level) modelling to the fNIRS data. Hierarchical models are able to account for within- and between-subject variability, making them less prone to bias than traditional summary measures (Boisgontier and Cheval, 2016; Gelman and Hill, 2006; Hesselmann, 2018). Modelling the data in this manner thus allows more accurate estimation of the neural differences between conditions (Galbraith et al., 2010). To this end, for each participant, we first removed outliers by removing any trials whose maximum absolute value exceeded 3 × *IQR* (interquartile range) of maximum values in that trial type (stop, go, ignore). We then normalised each trial by dividing it by the mean standard deviation (SD) of all trials for that participant per ROI. After normalisation, the local maximum of Hb concentration in each trial 2-7 seconds after the event trigger was extracted using custom Python code. This window was chosen given that changes in Hb concentration relevant to the presented stimulus tend to occur 3-6 s after stimulus presentation (Healey et al., 2024; Pinti et al., 2020). Analysis code for outlier removal, normalising, and the extraction of trialwise datapoints is shared at https://osf.io/ztb5m/.

### 2.5. Data Analysis

Trials are categorised as either *successful go* trials (two green arrows presented and the participant responds correctly), *failed go* trials (two green arrows presented and the participant makes some response other than a bimanual response), *failed stop* trials (a stop signal is presented but the participant responds with both hands, no hands, or with the wrong hand), *successful stop* trials (a stop signal is presented and the participant gives the correct, unimanual response), *successful ignore* trials (an ignore signal is presented and the participant gives a bimanual response), or *failed ignore* trials (an ignore signal is presented and the participant gives some response other than a bimanual button press). Trials in which the participant responded < 100 ms after go signal onset or slower than the stimulus duration of 2000 ms were excluded from the analysis (0.07% of all trials). Failed go and failed ignore trials were not included in the analyses due to their infrequency (participants responded correctly in 99.92% of go trials and 98.74% of ignore trials). Thus, unless otherwise specified, the terms ‘go trial’ and ‘ignore trial’ refer only to *correct* go and *correct* ignore trials, respectively.

For statistical analysis of the behavioural and EMG data, we used generalised linear mixed models (GLMMs), with either Gaussian distributions and log link functions, or gamma distributions with identity functions. Both approaches are effective in combating the skewness inherent in RT data (van der Linden, 2006; Lo and Andrews, 2015; Morís Fernández and Vadillo, 2020). The fNIRS data contained both positive and negative numbers and was less skewed (i.e., closer to a normal distribution), thus we used standard linear mixed models (LMMs) to assess them, which assume that the data are normally distributed.

In each case, multiple candidate models with varying random effects structures were fit (e.g., random intercepts only vs random intercepts plus random slopes). Candidate models were compared using Bayesian Information Criterion (BIC; Schwarz, 1978). The model with the lowest BIC was selected as the winning model. All candidate models and their relative fits (BICs) can be found in the Supplementary Materials. Models were implemented using the *GAMLj* package in jamovi v2.3 (Gallucci, 2019; Love et al., 2022, R Core Team 2021). The jamovi output for all analyses, including all candidate models, can be found at https://osf.io/ztb5m/.

#### 2.5.1. Behavioural analysis

For the behavioural analysis, in trials with bimanual responses, RT was quantified as the mean of the two button presses. In trials with a unimanual response (successful stop trials), RT was the timing of the single button press. Note that a unimanual button press was also possible in incorrect go and failed ignore trials, but these rarely occurred (0.05% and 0.15% of all trials, respectively), and were not included in the analysis.

In standard SSTs, there is no behavioural measure in successful stop trials as the participant, by definition, has cancelled their response. The stop-signal reaction time (SSRT) is often used an estimation of the latent stopping process (Logan and Cowan, 1984; Matzke et al., 2018; Verbruggen et al., 2013). In the current paradigm, the selective stopping means that there is an overt behavioural response in every trial, even in a successful stop trial (where there will be a unimanual button press from the unstopped hand). We thus assessed RT using a GLMM with trial type (go, failed stop, successful stop, and ignore) as a factor.

#### 2.5.2. EMG analysis

For the EMG analysis, we were primarily interested in examining partial responses. We thus focused on successful stop and successful ignore trials, assessing the effects of partial responses on other measures (i.e., RT and fNIRS measures). Partial responses could appear in one or both hands within a trial, so the RT and EMG measures were taken for each hand per trial, rather than being averaged across hands. We assessed the EMG measures in four different ways, as described below.

*RT* We assessed RT differences in trials with and without partial responses using a GLMM with trial type (successful stop, successful ignore) and partial response (present, absent) as factors. For this analysis, we took the RT of the uncued or unstopped hand in each trial. For a stop trial, we took the RT of the continuing hand, i.e., the one that *wasn’t* cued to stop. For an ignore trial, we took the RT of the hand for which there was no ignore signal. This was to ensure RT was comparable between stop and ignore trials.

*Cancel time (CT)* CT is the latency of the peak amplitude of the EMG signal in a partial response, relative to that trial’s SSD (see Figure 2). Previous work has used this measure as a trial-wise estimate of stopping latency (Jana et al., 2020; Weber et al., 2023), and it is considered a close electrophysiological correlate to SSRT (Raud et al., 2022; Salomoni et al., 2024). We assessed each hand individually per trial, as partial responses could occur in one or both hands in any trial. In this context, hands were classified as cued or uncued, depending on whether the stop or ignore signal appeared on that side. We assessed CT latency using a GLMM with trial type (stop, ignore) and hand (cued, uncued) as factors.

*Amplitude* We then assessed the amplitudes of response-generating bursts in successful stop and ignore trials (given by ω in Figure 2) with and without partial responses. As in the assessment of RT as a function of partial response presence (see above), we here took only the amplitude of the continuing hand, i.e., the left hand in a right ignore/stop trial, and the right hand in a left ignore/stop trial. We assessed amplitude using a GLMM with trial type (stop, ignore) and partial response (present, absent) as factors.

*Action reprogramming* Finally, we examined the speed of action reprogramming in stop and ignore trials with partial responses by assessing the latency between the peak of the partial response and the peak of the response-generating EMG burst (ω and γ in Figure 2, respectively). Again, in order to make the stop and ignore trials more comparative, we took the continuing (non-cued) hand (e.g., the left hand in a right ignore trial). We assessed these latencies using a GLMM with trial type (stop, ignore) as a factor.

#### 2.5.3. fNIRS analysis

As with the EMG analysis, we focused our analyses on successful stop and ignore trials, i.e., trials that could have partial responses. For each of the three ROIs, we assessed Hb concentration in stop and ignore trials with trial type (stop, ignore) and partial response (present, absent) as factors using an LMM.

## 3. Results

### 3.1. Behavioural results

Descriptive statistics summarising the behavioural data are reported in Table 2. The winning model had a significant main effect of trial type, *χ*^2^(3) = 6690.05, *p* < .001 (see Supplementary Materials for all candidate models). Holm-corrected post hoc tests found significant differences in mean RT between all trial types (all *p* < .001). Figure 4 shows the full RT distributions for each trial type in this analysis.

**Table 2.** Descriptive statistics of the behavioural data. Mean reaction time (RT) and standard deviation (SD) is given for all (successful) go trials, failed stops, successful stops, and (successful) ignores (SS = successful stop). Additionally, descriptive statistics for the SS and ignore trials are segmented based on the presence of partial EMG responses (PR). Failed ignore and failed go trials were excluded from the analysis due to their infrequency (participants made errors on 0.01% of go trials and 1.3% of ignore trials) and are thus not reported here.

| Trial type | Proportion | Mean RT (ms) | SD RT (ms) |
| --- | --- | --- | --- |
| Go |  | 498 | 170 |
| Failed stop |  | 432 | 132 |
| Successful stop (SS) |  | 641 | 170 |
| SS with a PR | 39.5% | 685 | 172 |
| SS without a PR | 60.5% | 611 | 167 |
| Ignore |  | 582 | 214 |
| Ignore with a PR | 26.1% | 724 | 213 |
| Ignore without a PR | 73.9% | 519 | 183 |

**Figure 4.**
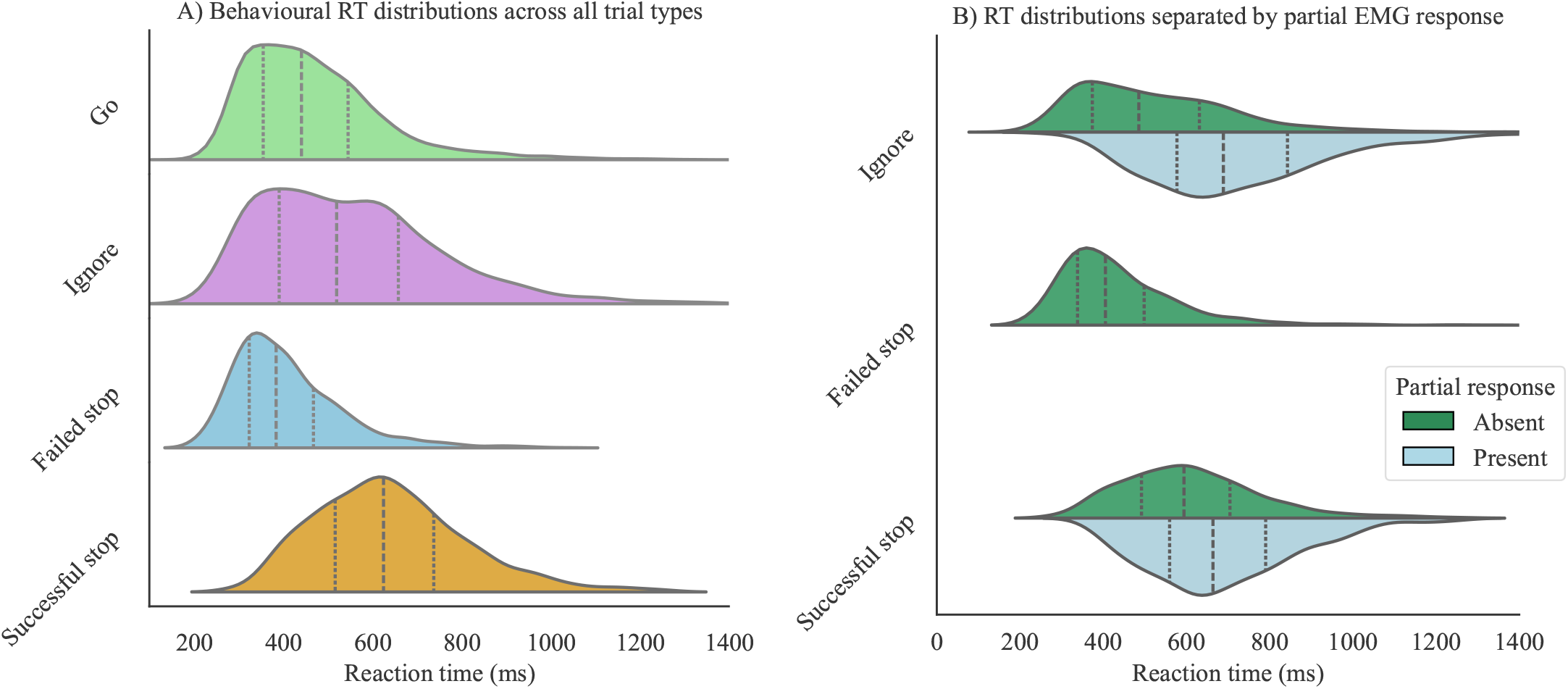
Reaction time (RT) distributions with quartiles for A) all trial types and B) stop and ignore trials, split by the presence of partial responses. Dotted lines represent the 25th and 75th percentiles; dashed lines represent the 50th. A) shows RT distributions for all four trial types (go trials, ignore trials, failed stops, and successful stops). There were significant differences between RT in all four trial types (all *p* < .001; see 3.1). B) shows the RT distributions for successful stop and ignore trials, split by the presence of partial EMG responses. All RT differences were significant (all *p* < .001; see 3.2.1). Note that failed stop trials were not included in the analysis for 3.2 as they do not contain partial responses, but are depicted here for comparative purposes.

**Figure 5.**
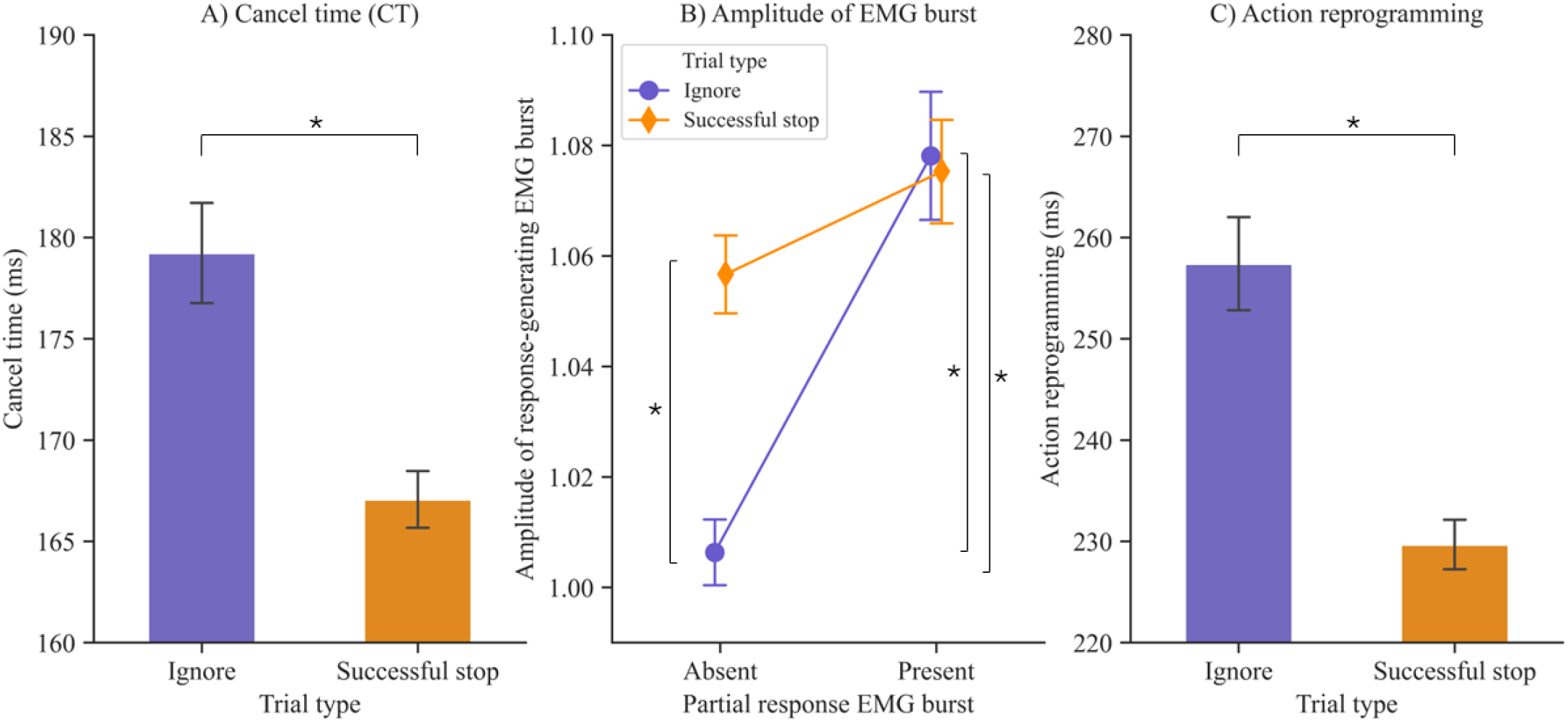
EMG results showing differences in cancel time (A), the amplitude of the response-generating burst (B), and action reprogramming (C) in stop and ignore trials. Error bars denote SE. Significant differences (*p* < .05) are denoted with *. Cancel time is the latency of the peak of a partial response relative to the stop-signal delay (SSD). Amplitude is relative to the mean amplitude in successful go trials. Action reprogramming is the duration between the peaks of a partial response and a response-generating RT burst.

### 3.2. EMG results (including RT results split by presence of partial responses)

Descriptive statistics summarising CT, amplitude, and action reprogramming in stop and ignore trials are reported in Table 3. Details of winning models and all candidate models are reported in the Supplementary Materials.

**Table 3.** Descriptive statistics for cancel time (CT), the amplitude of the response-generating RT burst, and action reprogramming for stop and ignore trials. Amplitude is relative to the mean amplitude in successful go trials. Ignore trials were significantly slower than stop trials for both CT and action reprogramming, both *p* < .001. The amplitude of the response-generating RT burst was significantly lower in ignore trials without a partial response (PR) than in the other conditions (all *p* ≤ 0.036).

| Measure | Trial type | Mean | SD |
| --- | --- | --- | --- |
| Cancel time (ms) | Stop | 167 | 57 |
|  | Ignore | 179 | 86 |
| Amplitude of response-generating RT burst | Stop with PR | 1.08 | 0.27 |
|  | Stop no PR | 1.06 | 0.25 |
|  | Ignore with PR | 1.08 | 0.28 |
|  | Ignore no PR | 1.01 | 0.25 |
| Action reprogramming (ms) | Stop | 230 | 71 |
|  | Ignore | 257 | 109 |

#### 3.2.1. Effect of partial response on reaction time (RT)

There was a significant interaction between partial response and trial type (*χ*^2^[1] = 459.35, *p* < .001), and significant main effects of both partial response (*χ*^2^[1] = 442.52, *p* < .001) and trial type (*χ*^2^[1] = 42.68, *p* < .001). Holm-corrected post hoc tests revealed there were significant differences in mean RT for all possible comparisons (all *p* < .001). In general, trials with a partial response had a longer RT than those without for both successful stop and ignore trials. Descriptive statistics are given in Table 2. See Figure 4 for RT distributions split by partial response presence.

#### 3.2.2. Cancel time (CT)

There was a statistically significant main effect of trial type (*χ*^2^[1] = 8.30, *p* = .004); CT was slower in ignore trials than in successful stop trials (Table 3). The interaction effect (trial type * hand) and main effect of hand were not significant (*χ*^2^[1] = 0.001, *p* = .975 and *χ*^2^[1] = 0.07, *p* = .795, respectively). See Figure 5.

#### 3.2.3. EMG amplitude in stop and ignore trials

There was a statistically significant interaction between trial type and partial response (*χ*^2^[1] = 10.49, *p* = .001). The interaction effect is depicted in Figure 5. Holm-corrected post hoc tests indicated that in ignore trials without a partial response, the amplitude of the subsequent response-generating burst was significantly lower than in stop trials without a partial response (*p* = .036), ignore trials with a partial response (*p* < .001), and stop trials with a partial response (*p* = .003). There was no significant difference in the response-generating amplitude of stop trials with and without partial responses (*p* = .403) or between ignore and stop trials with partial responses (*p* = .657). See Figure 5.

There was also statistically significant main effect of partial response (*χ*^2^[1] = 27.57, *p* < .001). The amplitude of the response-generating burst was higher in trials with a partial response (*M* = 1.08, *SD* = 0.27) than trials without one (*M* = 1.03, *SD* = 0.25). The main effect of trial type was not statistically significant (*χ*^2^[1] = 1.20, *p* = .273). See Figure 5.

#### 3.2.4. Action reprogramming in stop and ignore trials with partial responses

There was a significant effect of trial type (*χ*^2^[1] = 12.59, *p* < .001). The delay between partial responses and response-generating bursts was significantly longer in ignore trials (*M* = 257.43 ms, *SD* = 109.10) than in stop trials (*M* = 229.69 ms, *SD* = 71.13). See Figure 5.

### 3.3. fNIRS results

#### 3.3.1. preSMA

There were significant main effects of trial type and partial-response (*χ*^2^[1] = 5.78, *p* = .016 and *χ*^2^[1] = 3.97, *p* = .046, respectively). See Figure 6. Peak Hb concentration was significantly higher in successful stop trials (*M* = 0.94, *SD* = 1.59) than in ignore trials (*M* = 0.81, *SD* = 1.66). Peak Hb concentration was also significantly higher in trials *without* partial EMG responses (*M* = 0.92, *SD* = 1.60) than trials with them (*M* = 0.79, *SD* = 1.66). The interaction effect was not statistically significant (*χ*^2^[1] = 0.54, *p* = .464).

**Figure 6.**
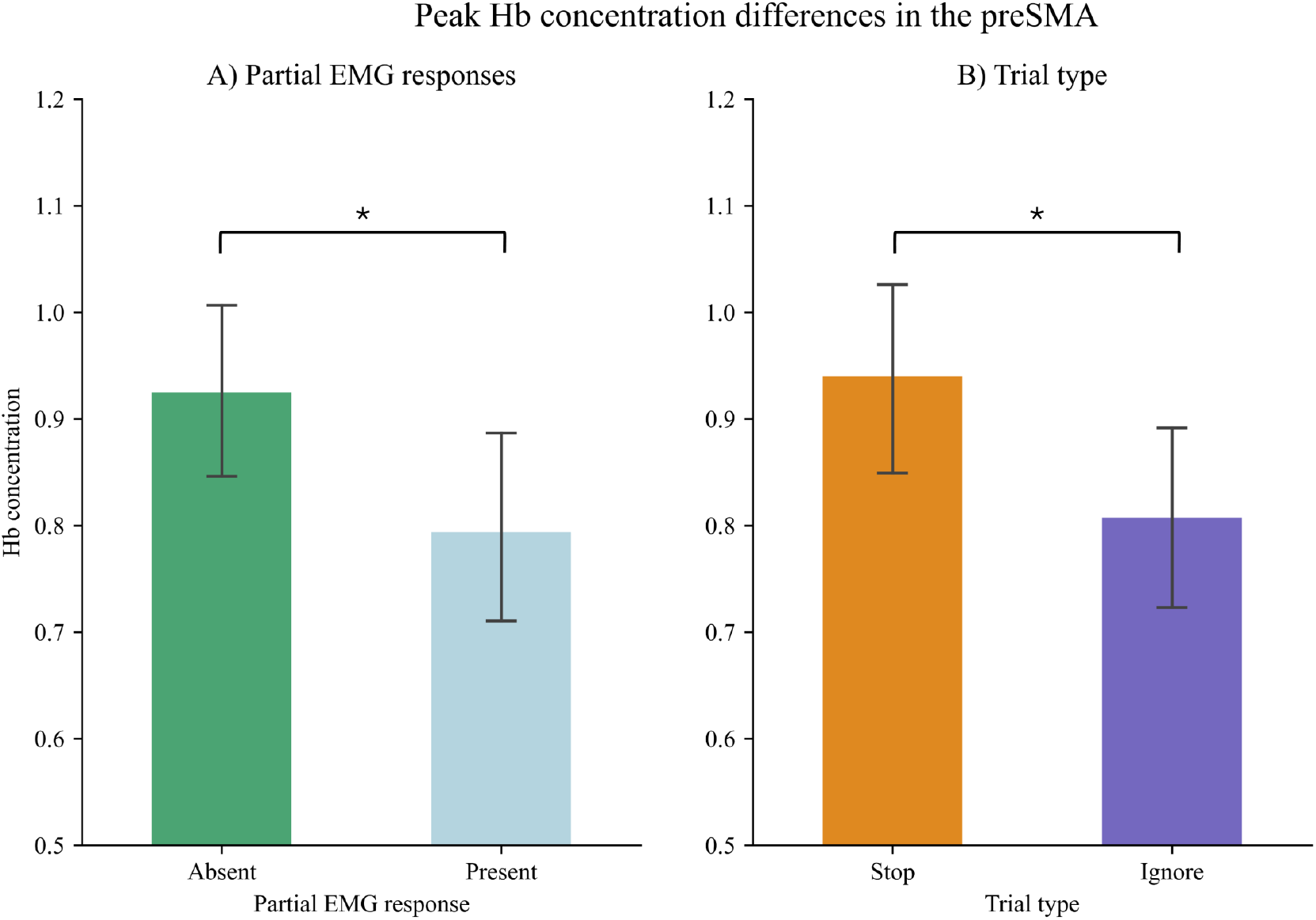
Peak oxygenated haemoglobin (Hb) concentration in the pre-supplementary motor area (preSMA). Increased Hb concentration indicates increased neural activity in that condition. There was significantly higher peak concentration in trials without a partial EMG response (A), and significantly higher peak concentration in successful stop trials (B).

#### 3.3.2. Right and left IFG

For the RIFG, neither of the main effects or interaction terms were statistically significant (trial type: *χ*^2^[1] = 1.90, *p* = .168; partial response: *χ*^2^[1] = 0.01, *p* = .928; interaction: *χ*^2^[1] = 0.30, *p* = .582).

Similarly, for the LIFG, none of the main effects or interaction terms were statistically significant (trial type: *χ*^2^[1] = 3.12, *p* = .077; partial response: *χ*^2^[1] = 0.19, *p* = .664; interaction: *χ*^2^[1] < 0.01, *p* = .983).

The interaction effects for the RIFG and LIFG are depicted in the Supplementary Materials, along with all candidate models.

## 4. Discussion

The preSMA and IFG have long been recognised for their crucial roles in action cancellation. Contemporary models of response inhibition, such as the PTC model, propose action cancellation occurs in two stages, with the IFG and preSMA underpinning the ‘pause’ and ‘cancel/retune’ processes, respectively (Diesburg and Wessel, 2021; Tatz et al., 2021). Previous research has also identified partial EMG responses in certain stopping scenarios, where a movement is initiated at the level of the effector muscle but no observable behavioural output ensues. The presence of partial EMG responses in some successful stop and ignore trials suggests that distinct processes may be at play in these instances (note that failed stop trials almost never have partial responses, hence here we focus on *successful* stop and ignore scenarios). Although partial responses are thought to reflect late-stage inhibition (Raud et al., 2022), their relationship with the regions that underpin action cancellation are yet to be explored. Here, we investigated potential neural correlates of these partial responses and their impact on overt behaviour.

### 4.1. Reaction time delay in both stop and ignore trials

Consistent with previous studies, RTs in successful stop trials were delayed compared to RTs in go trials, by approximately 140 ms (see Table 2 and Figure 4). This behavioural phenomenon is frequently to as the ‘stopping interference effect’ or ‘stopping delay’ (Gronau et al. 2024; Salomoni et al. 2023; for a review, see Wadsley et al., 2022). While this effect has been thought a consequence of the stopping process itself, it has primarily been investigated using standard SSTs, which involve only go and stop cues, meaning attentional cuing effects may have confounded this interpretation. To disentangle these mechanisms, some studies have, thus, incorporated attentional (ignore) cues in selective stopping paradigms.

Here, we observed that ignore trials also exhibit RT delays (≈ 80 ms delay in ignore trials; Table 2), consistent with previous works (Ko et al., 2015; Ko and Miller, 2013; Weber et al., 2023). Further insight into these RT delays can be gained by considering successful stop and ignore trials with and without partial responses (Figure 4). Ignore trials without a partial response are nearly 100 ms faster than successful stop trials without a partial response (Table 2). Notably, successful ignore trials are trials where the ignore signal is presented and the participant makes a bimanual response. They therefore contain trials where the response would have been executed prior to perceiving the ignore signal, as the correct response in this instance is the prepotent response. Conversely, in stop trials, if the participant makes a bimanual response before the stop signal is perceived, then this will count as a failed trial. Consequently, comparing trials with partial responses may be more informative, as these represent instances where the prepotent response has been modified or paused in some way (and the stop/ignore signal has been processed). In this context, we see that ignore trials with partial responses are actually ≈ 40 ms *slower* than successful stop trials with a partial response (Table 2). This effect is consistent in other EMG measures (CT and action reprogramming), which are discussed below.

### 4.2. Cancel time (CT) and action reprogramming are slower in ignores than in stop trials

Trials with partial responses likely represent instances where a participant has initiated a movement that is then modified or reinitiated, depending on contextual requirements (Wadsley and Greenhouse, 2024). We assessed *CT* (the latency of the peak of a partial response, relative to SSD) and *action reprogramming* (the time between the peaks of the partial response and the RT-generating EMG burst; Weber et al. 2024a^1^), and found both of these measures to be delayed in ignore trials compared to stop trials.

Longer CT in ignore trials relative to stop trials (Table 3) is inconsistent with previous work, which has found no evidence of CT differences between stop and ignore trials (Tatz et al. 2021; Weber et al. 2024b). Notably, there are some procedural differences between these prior studies and the current work with respect to outlier removal. Both papers used a relatively conservative outlier approach; Tatz and colleagues removed EMG trials identified according to Cook’s distance (using a threshold of 4/*N*), while Weber and colleagues removed any trials where CT was more than 2 SDs above the mean. Here, we chose not to truncate the dataset by removal of upper-limit trials, instead fitting a series of candidate models with varying distributions, which can accommodate the skews frequently observed in RT data (for discussions on RT distributions and outlier removal, see Heathcote et al., 1991; Miller, 1991; Ulrich and Miller, 1994). While it seems logical to assume that the onset of partial responses in stop and ignore trials reflects the same underlying physiological process (as has been previously argued; Tatz et al., 2021; Weber et al., 2024a), it is also clear that ignore trials have a broader distribution than stop trials, specifically with a wider upper tail.

We assessed the speed of action reprogramming in stop and ignore trials by measuring the delay between the onsets of the peaks of partial responses and the peaks of the response-generating RT burst. We found that the speed of action reprogramming was slower for ignore trials than stop trials by approximately 30 ms (Table 3), consistent with previous findings (Weber et al., 2024a). This is also evident in the behavioural RT, where ignore trials with partial responses are slower than stop trials with partial responses, as discussed above (see Figure 5 and Table 3).

The delay in CT and action reprogramming in ignore trials has interesting implications for models of action cancellation. Under the PTC model, the pause process is thought to be triggered non-specifically by a stop or ignore signal, delaying the ongoing action but not modifying it. Following the pause, the specific imperative to stop associated with the stop signal is thought to trigger the cancel/retune process, which modifies the ongoing action and enables action cancellation. Action in ignore trials is thus conceptualised as the original bimanual response that has been delayed (due to the pause process). Thus, it seems counterintuitive that these trials would be delayed relative to instances where an action has been paused *and* modified, to result in a reprogrammed response (i.e., in a stop trial).

Whilst the PTC model proposes that selective stopping entails the modification of an *ongoing* movement, under cognitive modelling approaches such as the Activation Threshold Model (ATM; MacDonald et al., 2017) and the Simultaneously Inhibit and Start (SIS) model (Gronau et al., 2024), actions are not proposed to be modified *in situ*, but rather entirely inhibited before a new action is initiated. Such approaches have not, as of yet, considered RT delays in non-stopping contexts, instead focusing on stop versus go scenarios. These frameworks may need to be generalised in the future to incorporate a greater range of action modification scenarios, where they may be well-suited to answering how the onset of an infrequent stimulus without an imperative to stop may be managed. Speculatively, in this context, the behavioural responses in ignore trials could be considered the result of the original bimanual process, triggered by go onset (as is the case in the PTC model), or that of a subsequent process, initiated later in the trial in response to the ignore stimulus.

### 4.3. Increased amplitude of response-generating EMG bursts in trials with partial responses

We assessed how partial responses in stop and ignore trials affected the amplitudes of the subsequent response-generating RT bursts (ω in Figure 2). We found that ignore trials with partial responses and successful stop trials (with or without partial responses) all had significantly higher amplitudes than ignore trials without partial responses. In Figure 5, it can be seen that the amplitude of the ignore trials without partial responses was very close to 1, i.e., close to the amplitude of standard bimanual go trials. Ignore trials without partial responses are likely to represent something very close to these standard go trials, i.e., where the participant makes the prepotent bimanual response. Conversely, ignore trials *with* partial responses and successful stop trials (with or without partial responses) represent instances where response modification has taken place. Either the bimanual response has been preceded by a partial response (in an ignore trial) or the action has been modified/cancelled and a unimanual response executed instead (in a stop trial). This is consistent with both the ATM and SIS models, which propose that the re-initiated response needs to be stronger than the original, bimanual response in order to overcome the ongoing inhibition in corticomotor pathways (Gronau et al., 2024; MacDonald et al., 2017)

### 4.4. IFG and preSMA activity in stop trials compared to ignore trials

To our knowledge, this is the first study to use fNIRS to investigate the neural correlates of action cancellation, with previous experimental work predominantly employing fMRI and brain stimulation techniques. In order to isolate the neural correlates of stopping and attentional capture, we examined neural activity in the preSMA, RIFG, and LIFG in (successful) stop and ignore trials, finding increased neural activity in (successful) stop compared to ignore trials in the preSMA (Figure 6), but not in the left or right IFG.

The roles of these regions in action cancellation have been extensively investigated, primarily by means of standard (non-selective) SSTs, where brain activation (measured via the BOLD signal) is contrasted between go trials, successful stops, and failed stops. Increased preSMA and bilateral IFG activation have consistently been observed in failed and successful stop trials compared to go trials (see e.g., Aron and Poldrack, 2006; Boehler et al., 2011; Chao et al., 2009; Messel et al., 2019). Disruption or facilitation of these regions using brain stimulation has found subsequent impairment or improvement in SST performance (Chen et al., 2009; Kohl et al., 2019; Lee et al., 2016; Watanabe et al., 2015), although this is not always observed (Friehs et al., 2023; Obeso et al., 2017).

As discussed above, the infrequent nature of the stop signal in standard SSTs means these trials are likely capturing both action cancellation and attentional orienting effects (Banich and Depue, 2015), which has led some authors to question the extent to which these activation patterns can be attributed solely to action cancellation (Hampshire, 2015; Sharp et al., 2010). To control for these attentional effects, some studies have instead contrasted (successful) stop trials with *ignore* trials. These have yielded markedly different effects, with clusters of activation found in the preSMA and little to no activation found in the (right) IFG (Sharp et al., 2010). The current study, using fNIRS, reports evidence consistent with this fMRI data, with significant increases in activity in the preSMA in stop trials compared to ignore trials, and no significant differences found between stop and ignore trials in the right or left IFG. IFG activation following presentation of unexpected or infrequent stimuli has been observed in a variety of paradigms (Boehler et al., 2011; Chatham et al., 2012; Erika-Florence et al., 2014; Sebastian et al., 2020), suggesting its role in an action cancellation contexts may indeed be that of attentional orienting, i.e., not specific to action cancellation itself. This is also consistent with the PTC framework; the IFG is proposed to trigger the pause process in response to *any* infrequent or unexpected stimuli, meaning it would equally active in stop and ignore conditions. Conversely, the preSMA is thought to initiate the cancel/retune process when action cancellation is required, meaning preSMA activity would be expected in stop but not ignore conditions, as was observed here.

### 4.5. IFG and preSMA activity in trials with partial responses

The identification of partial EMG responses has become an increasingly-important component of action cancellation research. As well as providing an objective measurement of the dynamic muscular changes that occur during a trial, EMG allows measurement of latent processes not identifiable by standard behavioural measures (Raud et al., 2022; Thunberg et al., 2024). Here, we present an investigation of the neural correlates of these partial responses. We did not find any significant differences in IFG activation between trials with and without partial EMG responses, but found significantly more activity in the preSMA in trials *without* partial responses (Figure 6).

Wadsley and Greenhouse (2024) recently proposed that the presence of partial EMG responses in SSTs may be a way to delineate between trials with distinct underlying processes. Particularly, they suggest that successful stop trials without partial responses may pertain to trials where action *withholding* took place, while those with partial responses are instances of true action cancellation. Action withholding is the complete withholding of a motor response, normally associated with tasks such as go/no-go tasks, in which ‘no-go’ stimuli are infrequently presented in place of the normal ‘go’ stimuli at the same (predictable) time point. Participants thus wait to initiate their action until stimulus onset, requiring inhibition of a prepotent response, but *not* after another movement has been initiated. Differences in neural correlates between action withholding and action cancellation have previously been examined by Sebastian et al. (2013), who used a novel response inhibition task (the Hybrid Response Inhibition Task) with action cancellation, action withholding, and interference inhibition components, allowing comparisons of these mechanisms in the same sample. They found BOLD activity overlapped in a number of regions for all three components in their task (preSMA, right inferior frontal cortex [IFC]^2^, right inferior frontal junction, and bilateral parietal regions). Notably, they found there was more activity in the bilateral IFC, preSMA, right striatum, and left parietal lobule for action cancellation than for action withholding.

If successful stop trials without partial responses do, as proposed, represent instances of action withholding rather than action cancellation, increased activity in the ROIs in trials with partial responses would have been predicted, based on the results highlighted above from previous studies. Instead, we found no evidence of differences between trials with and without partial responses for the IFG, and actually found increased preSMA activity in trials *without* partial responses. There are a number of considerations to be made here. First, in contrasting instances of action cancellation and action withholding, Sebastian et al. (2013) contrasted successful stop trials with no-go trials. EMG was not collected in their experiment, but the successful stop trials presumably contained trials with and without partial responses.

Thus, this comparison may have actually contrasted action withholding (no-go) trials against trials with some instances of action withholding and some of action cancellation (successful stop trials). Nonetheless, the opposite effect observed here (increased activation in trials *without* partial responses) is unexpected under this interpretation, and indicates that these trials are unlikely to be instances of action withholding.

The preSMA and IFG in action cancellation are generally thought to be involved in the *initiation* of action cancellation mechanisms. Following these initiation points, the pathways are thought to involve a broad range of basal ganglia regions (striatum, globus pallidus externa and interna, and the STN), as well as the thalamus and M1, without rerouting back to the preSMA and IFG (Graybiel, 2000; Rocha et al., 2023; Schroll and Hamker, 2013). It is possible that action cancellation signals sent by either region may interrupt neural activity anywhere along this pathway, e.g., by increasing the thalamus’ inhibitory drive on M1 before the go command has been issued. In such cases, no partial response would be observed in that trial, as the motor command would not have reached the neuromuscular junction, but action cancellation mechanisms would still be responsible for the successful stop. Thus, action cancellation mechanisms may still underpin successful stop trials without partial responses.

However, even if action cancellation mechanisms underpin successful stopping, this does not explain the increase in preSMA activity in trials without partial responses. Here, proactive inhibition mechanisms are worth considering. Proactive inhibition a widely-used term that has been applied in a broad range of experimental paradigms, but generally, is considered a preparatory process that works to facilitate action cancellation, should it be required (for reviews, see Meyer and Bucci, 2016; van den Wildenberg et al., 2022). Importantly, it is thought to influence the likelihood that a response can be cancelled through pre-emptive modulation of corticospinal excitability or slowing of action initiation (Bundt and Huster, 2024; Puri et al., 2018). It is thought to be more active in complex stopping scenarios, e.g., where participants are given information about the probability of the upcoming stop trial being a stop trial (Hu et al., 2019; Messel et al., 2019; Meyer and Bucci, 2016; Vink et al., 2015). Proactive inhibition can be contrasted with reactive inhibition, when the cessation of the motor process is initiated entirely *after* onset of the stop stimulus. Reactive inhibition has largely been the focus of action cancellation research, where participants are not given information about the upcoming trial and cannot prepare appropriately.

Reactive and proactive inhibition are not considered mutually exclusive; indeed, it has been proposed that *both* may be required for effective inhibitory control (Criaud et al., 2012). The preSMA has been associated with both reactive and proactive inhibition (Criaud et al., 2012; Kenemans, 2015; Meyer and Bucci, 2016; Obeso et al., 2013). Evidence from non-human primates suggests that different neural populations within the preSMA may modulate these two types of inhibition (Chen et al., 2010). It has also been suggested that the preSMA may drive proactive *and* reactive inhibition at different time points in humans (proactive followed by reactive; Zhu et al., 2024). Speculatively, the increased preSMA activity in trials without partial responses observed in the current study may be reflective of *both* proactive and reactive inhibitory processes at work, whereas trials with partial responses may represent instances of reactive inhibition alone. Notably, proactive inhibition has been associated with a much wider range of neural correlates than reactive inhibition. Here, we investigated only two ROIs; future work may be able to examine differences in partial EMG responses in a broader range of regions, perhaps investigating regions that have been associated *solely* with proactive inhibition, such as regions in the superior parietal lobe (van Belle et al., 2014).

### 4.6. Limitations

In the current study, we assessed the ROIs individually, running models on their changes in haemoglobin concentration and assessing whether they differed between trial conditions or in instances with and without partial responses. Given that these regions are different sizes with differing vasculature, and that the preSMA is further from the skull than the IFG, we chose to assess these regions individually (in separate models) due to fundamental differences in their baseline activity. We maintain that direct comparisons here are limited in meaning. Nonetheless, this means that we cannot make statements about relative amounts of Hb concentration (e.g., we cannot state if the IFG is *more* associated with ignore signals than the preSMA; Nieuwenhuis et al., 2011).

### 4.7. Conclusion

The preSMA and IFG have long been associated with action cancellation. Two-stage models such as the PTC model hypothesise that these regions play quite distinct roles in coordinating the termination of action. Partial EMG responses have also been investigated extensively in action cancellation paradigms, although their neural correlates are unknown. Here, we assessed the role of the preSMA and IFG in action cancellation, attentional orienting, and partial EMG responses using a novel fNIRS paradigm. We found that the preSMA was uniquely involved in action cancellation while the IFG appeared equally involved in action cancellation and attentional capture, as would be predicted by the PTC model. We found greater preSMA activity in trials without partial EMG responses, and propose that this may result from increased proactive inhibition in these trials, coupled with the reactive inhibition mechanisms involved in trials with and without partial responses.

## Acknowledgements

The authors express their gratitude to Niek Stevenson and Simon Weber for insightful discussions regarding data analysis and interpretation, and to Rebecca Healey for her invaluable assistance in data collection and preprocessing.

## CRediT authorship contribution statement

**Sarah A Kemp:** Conceptualisation, Data curation, Formal analysis, Investigation, Methodology, Project administration, Software, Validation, Visualisation, Writing – original draft, Writing – review & editing.. **Sauro Salomoni:** Data curation, Methodology, Software, Validation, Writing – review & editing.. **Pierre-Louis Bazin:** Methodology, Software, Supervision, Validation, Writing – review & editing.. **Luke Pash:** Data curation, Investigation, Writing – original draft, Writing – review & editing.. **Rebecca J St George:** Data curation, Funding acquisition, Resources, Supervision, Validation, Writing – review & editing.. **Mark R Hinder:** Conceptualisation, Funding acquisition, Resources, Supervision, Writing – review & editing..

## Footnotes

1 Note that Weber and colleagues defined action reprogramming to be the time between the peak of the partial response and the *onset* of the RT-generating burst, rather than the time between the two peaks. The width of the response-generating EMG burst is consistent between successful stops and standard go trials (Fisher et al., 2024), so these two approaches can be conceptualised in the same manner.

2 Note that the IFC is a larger region than the IFG, including both the IFG and the frontal operculum (Frühholz and Grandjean, 2013).

## References

Aron, A.R., Poldrack, R.A., 2006. Cortical and subcortical contributions to stop signal response inhibition: Role of the subthalamic nucleus. Journal of Neuroscience 26, 2424–2433. URL: https://www.jneurosci. org/content/26/9/2424, doi:10.1523/JNEUROSCI.4682-05.2006.

Aron, A.R., Verbruggen, F., 2008. Stop the presses: Dissociating a selective from a global mechanism for stopping. Psychological Science 19, 1146–1153. URL: http://journals.sagepub.com/doi/10.1111/j.1467-9280.2008.02216.x, doi:10.1111/j.1467-9280.2008.02216.x.

Atsma, J., Maij, F., Gu, C., Medendorp, W.P., Corneil, B.D., 2018. Active braking of whole-arm reaching movements provides single-trial neuro-muscular measures of movement cancellation. Journal of Neuroscience 38, 4367–4382. doi:10.1523/JNEUROSCI.1745-17.2018.

Banich, M.T., Depue, B.E., 2015. Recent advances in understanding neural systems that support inhibitory control. Current Opinion in Behavioral Sciences 1, 17–22. URL: https://linkinghub.elsevier.com/retrieve/pii/S2352154614000084, doi:10.1016/j.cobeha.2014.07.006.

van Belle, J., Vink, M., Durston, S., Zandbelt, B.B., 2014. Common and unique neural networks for proactive and reactive response inhibition revealed by independent component analysis of functional MRI data. NeuroImage 103, 65–74. URL: http://www.sciencedirect.com/science/article/pii/S1053811914007563, doi:10.1016/j.neuroimage.2014.09.014.

Benis, D., David, O., Piallat, B., Kibleur, A., Goetz, L., Bhattacharjee, M., Fraix, V., Seigneuret, E., Krack, P., Chabardès, S., Bastin, J., 2016. Response inhibition rapidly increases single-neuron responses in the subthalamic nucleus of patients with Parkinson’s disease. Cortex 84, 111–123. doi:10.1016/j.cortex.2016.09.006.

Bingham, C.S., Petersen, M.V., Parent, M., McIntyre, C.C., 2023. Evolving characterization of the human hyperdirect pathway. Brain Structure and Function 228, 353–365. URL: https://doi.org/10.1007/s00429-023-02610-5, doi:10.1007/s00429-023-02610-5.

Boehler, C.N., Appelbaum, L.G., Krebs, R.M., Chen, L.C., Woldorff, M.G., 2011. The role of stimulus salience and attentional capture across the neural hierarchy in a stop-signal task. PLOS ONE 6, e26386. URL: https://journals.plos.org/plosone/article?id=10.1371/journal.pone.0026386, doi:10.1371/journal.pone.0026386. publisher: Public Li-brary of Science.

Boehler, C.N., Appelbaum, L.G., Krebs, R.M., Hopf, J.M., Woldorff, M.G., 2010. Pinning down response inhibition in the brain — Conjunction analyses of the Stop-signal task. NeuroImage 52, 1621–1632. URL: http://www.sciencedirect.com/science/article/pii/S105381191000707X, doi:10.1016/j.neuroimage.2010.04.276.

Boen, R., Raud, L., Huster, R.J., 2022. Inhibitory control and the structural parcelation of the right inferior frontal gyrus. Frontiers in Human Neuroscience 16, 787079. URL: https://www.ncbi.nlm.nih.gov/pmc/articles/PMC8907402/, doi:10.3389/fnhum.2022.787079.

Boisgontier, M.P., Cheval, B., 2016. The ANOVA to mixed model transition. Neuroscience & Biobehavioral Reviews 68, 1004–1005. URL: http://www.sciencedirect.com/science/article/pii/S0149763416301634, doi:10.1016/j.neubiorev.2016.05.034.

Borgomaneri, S., Serio, G., Battaglia, S., 2020. Please, don’t do it! Fifteen years of progress of non-invasive brain stimulation in action inhibition. Cortex 132, 404–422. URL: https://linkinghub.elsevier.com/retrieve/pii/S0010945220303300, doi:10.1016/j.cortex.2020.09.002.

Borst, B., Jovanovic, T., House, S.L., Bruce, S.E., Harnett, N.G., Roeckner, A.R., Ely, T.D., Lebois, L.A.M., Young, D., Beaudoin, F.L., An, X., Neylan, T.C., Clifford, G.D., Linnstaedt, S.D., Germine, L.T., Bollen, K.A., Rauch, S.L., Haran, J.P., Storrow, A.B., Lewandowski, C., Musey, P.I., Hendry, P.L., Sheikh, S., Jones, C.W., Punches, B.E., Hudak, L.A., Pascual, J.L., Seamon, M.J., Datner, E.M., Pearson, C., Peak, D.A., Domeier, R.M., Rathlev, N.K., O’Neil, B.J., Sergot, P., Sanchez, L.D., Harte, S.E., Koenen, K.C., Kessler, R.C., McLean, S.A., Ressler, K.J., Stevens, J.S., van Rooij, S.J.H., 2024. Sex differences in response inhibition-related neural predictors of PTSD in recent trauma-exposed civilians. Biological Psychiatry: Cognitive Neuroscience and Neu-roimaging URL: https://www.sciencedirect.com/science/article/pii/S2451902224000806, doi:10.1016/j.bpsc.2024.03.002.

Brigadoi, S., Cooper, R.J., 2015. How short is short? Optimum source–detector distance for short-separation channels in functional near-infrared spectroscopy. Neurophotonics 2, 025005. URL: https://www.ncbi.nlm.nih.gov/pmc/articles/PMC4478880/, doi:10.1117/1.NPh.2.2.025005.

Bundt, C., Huster, R.J., 2024. Corticospinal excitability reductions during action preparation and action stopping in humans: Different sides of the same inhibitory coin? Neuropsychologia 195, 108799. URL: https://linkinghub.elsevier.com/retrieve/pii/S0028393224000149, doi:10.1016/j.neuropsychologia.2024.108799.

Chao, H.H., Luo, X., Chang, J.L., Li, C.s.R., 2009. Activation of the pre-supplementary motor area but not inferior prefrontal cortex in association with short stop signal reaction time – an intra-subject analysis. BMC Neuroscience 10, 75. URL: https://www.ncbi.nlm.nih.gov/pmc/articles/PMC2719646/, doi:10.1186/1471-2202-10-75.

Chatham, C.H., Claus, E.D., Kim, A., Curran, T., Banich, M.T., Munakata, Y., 2012. Cognitive control reflects context monitoring, not motoric stopping, in response inhibition. PLOS ONE 7, e31546. URL: https://journals.plos.org/plosone/article?id=10.1371/journal.pone.0031546, doi:10.1371/journal.pone.0031546.

Chen, C.Y., Muggleton, N.G., Tzeng, O.J.L., Hung, D.L., Juan, C.H., 2009. Control of prepotent responses by the superior medial frontal cortex. NeuroImage 44, 537–545. doi:10.1016/j.neuroimage.2008.09.005.

Chen, X., Scangos, K.W., Stuphorn, V., 2010. Supplementary motor area exerts proactive and reactive control of arm movements. The Journal of Neuroscience 30, 14657–14675. doi:10.1523/JNEUROSCI.2669-10.2010.

Coudé, D., Parent, A., Parent, M., 2018. Single-axon tracing of the corticosubthalamic hyperdirect pathway in primates. Brain Structure and Function 223, 3959–3973. URL: https://doi.org/10.1007/s00429-018-1726-x, doi:10.1007/s00429-018-1726-x.

Coxon, J.P., Stinear, C.M., Byblow, W.D., 2007. Selective inhibition of movement. Journal of Neurophysiology 97, 2480–2489. URL: http://journals.physiology.org/doi/full/10.1152/jn.01284.2006, doi:10.1152/jn.01284.2006.

Criaud, M., Wardak, C., Ben Hamed, S., Ballanger, B., Boulinguez, P., 2012. Proactive inhibitory control of response as the default state of executive control. Frontiers in Psychology 3. URL: https://www.frontiersin.org/journals/psychology/articles/10.3389/fpsyg.2012.00059/full, doi:10.3389/fpsyg.2012.00059. publisher: Frontiers.

Diesburg, D.A., Wessel, J.R., 2021. The Pause-then-Cancel model of human action-stopping: Theoretical considerations and empirical evidence. Neuroscience & Biobehavioral Reviews 129, 17–34. URL: https://linkinghub.elsevier.com/retrieve/pii/S0149763421003183, doi:10.1016/j.neubiorev.2021.07.019.

Eggebrecht, A.T., White, B.R., Ferradal, S.L., Chen, C., Zhan, Y., Snyder, A.Z., Dehghani, H., Culver, J.P., 2012. A quantitative spatial comparison of high-density diffuse optical tomography and fMRI cortical mapping. NeuroImage 61, 1120–1128. URL: https://www.sciencedirect.com/science/article/pii/S1053811912001516, doi:10.1016/j.neuroimage.2012.01.124.

Erika-Florence, M., Leech, R., Hampshire, A., 2014. A functional network perspective on response inhibition and attentional control. Nature Communications 5, 4073. URL: https://www.nature.com/articles/ncomms5073, doi:10.1038/ncomms5073. publisher: Nature Publish-ing Group.

Fisher, M., Trinh, H., O’Neill, J., Greenhouse, I., 2024. Early rise and persistent inhibition of electromyography during failed stopping. Journal of Cognitive Neuroscience 36, 1412–1426. URL: https://doi.org/10.1162/jocn_a_02174, doi:10.1162/jocn_a_02174.

Forstmann, B.U., Jahfari, S., Scholte, H.S., Wolfensteller, U., van den Wildenberg, W.P.M., Ridderinkhof, K.R., 2008. Function and structure of the right inferior frontal cortex predict individual differences in response inhibition: a model-based approach. Journal of Neuroscience 28, 9790–9796. doi:10.1523/JNEUROSCI.1465-08.2008.

Frank, M.J., 2006. Hold your horses: a dynamic computational role for the subthalamic nucleus in decision making. Neural Networks: The Official Journal of the International Neural Network Society 19, 1120–1136. doi:10.1016/j.neunet.2006.03.006.

Friehs, M.A., Hartwigsen, G., Matzke, D., Donzallaz, M.C., Siodmiak, J., Numssen, O., Frings, C., 2023. Effects of 1 Hz offline TMS on performance in the stop-signal game. preprint. Open Science Framework. URL: https://osf.io/a3ths, doi:10.31219/osf.io/a3ths.

Frühholz, S., Grandjean, D., 2013. Processing of emotional vocalizations in bilateral inferior frontal cortex. Neuroscience & Biobehavioral Reviews 37, 2847–2855. URL: https://www.sciencedirect.com/science/article/pii/S0149763413002315, doi:10.1016/j.neubiorev.2013.10.007.

Galbraith, S., Daniel, J.A., Vissel, B., 2010. A study of clustered data and approaches to Its analysis. Journal of Neuroscience 30, 10601–10608. URL: https://www.ncbi.nlm.nih.gov/pmc/articles/PMC6634702/, doi:10.1523/JNEUROSCI.0362-10.2010.

Gallucci, M., 2019. GAMLj: General Analyses for the Linear Model in Jamovi. URL: https://gamlj.github.io/.

Gavazzi, G., Lenge, M., Bartoli, E., Bianchi, A., Agovi, H., Mugnai, F., Guerrini, R., Giordano, F., Viggiano, M., Mascalchi, M., 2019. Left inferior frontal cortex can compensate the inhibitory functions of right inferior frontal cortex and pre-supplementary motor area. Journal of Neurophysiology 13, 503–508. URL: http://www.scopus.com/record/display.uri?eid=2-s2.0-85052817270&origin=resultslist&sort=plf-f&cite=2-s2.0-84897041787&src=s&nlo=&nlr=&nls=&imp=t&sid=e7f0e21cc14c6e72352d2ff0bf752957&sot=cite&sdt=cl&cluster=scosubjabbr%2c%22NEUR%22%2ct&sl=0&relpos=278&citeCnt=5&searchTerm=, doi:10.1111/jnp.12170.

Gelman, A., Hill, J., 2006. Data Analysis Using Regression and Multi-level/Hierarchical Models. Cambridge University Press, Cambridge.

Graybiel, A.M., 2000. The basal ganglia. Current Biology 10, R509–R511. URL: https://www.sciencedirect.com/science/article/pii/S0960982200005935, doi:10.1016/S0960-9822(00)00593-5.

Gronau, Q.F., Hinder, M.R., Salomoni, S.E., Matzke, D., Heathcote, A., 2024. A unified account of simple and response-selective inhibition. Cognitive Psychology 149, 101628. URL: https://www.sciencedirect.com/science/article/pii/S0010028523000865, doi:10.1016/j.cogpsych.2023.101628.

Hampshire, A., 2015. Putting the brakes on inhibitory models of frontal lobe function. NeuroImage 113, 340–355. URL: http://www.sciencedirect.com/science/article/pii/S1053811915002487, doi:10.1016/j.neuroimage.2015.03.053.

Hannah, R., Jana, S., Muralidharan, V., 2022. Does action-stopping involve separate pause and cancel processes? A view from premotor cortex. Cortex 152, 157–159. URL: https://www.sciencedirect.com/science/article/pii/S001094522100246X, doi:10.1016/j.cortex.2021.06.015.

Healey, R., Goldsworthy, M., Salomoni, S., Weber, S., Kemp, S., Hinder, M.R., St George, R.J., 2024. Impaired motor inhibition during perceptual inhibition in older, but not younger adults: a psychophysiological study. Scientific Reports 14, 2023. URL: https://www.nature.com/articles/s41598-024-52269-z, doi:10.1038/s41598-024-52269-z.

Heathcote, A., Popiel, S.J., Mewhort, D.J., 1991. Analysis of response time distributions: An example using the Stroop task. Psychological Bulletin 109, 340–347. URL: https://doi.apa.org/doi/10.1037/0033-2909.109.2.340, doi:10.1037/0033-2909.109.2.340.

Heeger, D.J., Ress, D., 2002. What does fMRI tell us about neuronal activity? Nature Reviews Neuroscience 3, 142–151. URL: https://www.nature.com/articles/nrn730, doi:10.1038/nrn730.

Hesselmann, G., 2018. Applying linear mixed effects models in within-participant designs with subjective trial-based assessments of awareness - a caveat. Frontiers in Psychology 9. URL: https://www.frontiersin.org/journals/psychology/articles/10.3389/fpsyg.2018.00788/full, doi:10.3389/fpsyg.2018.00788. publisher: Frontiers.

Hiroshima, S., Anei, R., Murakami, N., Kamada, K., 2014. Functional localization of the supplementary motor area. Neurologia medico-chirurgica 54, 511–520. URL: https://www.ncbi.nlm.nih.gov/pmc/articles/PMC4533467/, doi:10.2176/nmc.oa.2012-0321.

Hodges, P.W., Bui, B.H., 1996. A comparison of computer-based methods for the determination of onset of muscle contraction using electromyography. Electroencephalography and Clinical Neurophysiology/Electromyography and Motor Control 101, 511–519. URL: https://www.sciencedirect.com/science/article/pii/S0921884X96951905, doi:10.1016/S0921-884X(96)95190-5.

Hu, S., Job, M., Jenks, S., Chao, H., Li, C.S., 2019. Imaging the effects of age on proactive control in healthy adults. Brain Imaging and Behavior 13, 1526–1537. doi:10.1007/s11682-019-00103-w.

Hughes, M.E., Fulham, W.R., Johnston, P.J., Michie, P.T., 2012. Stop-signal response inhibition in schizophrenia: Behavioural, event-related potential and functional neuroimaging data. Biological Psychology 89, 220–231. URL: http://www.sciencedirect.com/science/article/pii/S0301051111002651, doi:10.1016/j.biopsycho.2011.10.013.

Huppert, T.J., Diamond, S.G., Franceschini, M.A., Boas, D.A., 2009. HomER: a review of time-series analysis methods for near-infrared spectroscopy of the brain. Applied Optics 48, D280–D298. URL: https://opg.optica.org/ao/abstract.cfm?uri=ao-48-10-D280, doi:10.1364/AO.48.00D280.

Jahfari, S., Waldorp, L., Wildenberg, W.P.M.v.d., Scholte, H.S., Ridderinkhof, K.R., Forstmann, B.U., 2011. Effective connectivity reveals important roles for both the hyperdirect (fronto-subthalamic) and the indirect (fronto-striatal-pallidal) fronto-basal ganglia pathways during response inhibition. Journal of Neuroscience 31, 6891–6899. URL: https://www.jneurosci.org/content/31/18/6891, doi:10.1523/JNEUROSCI.5253-10.2011. publisher: Society for Neuroscience Section: Articles.

Jana, S., Hannah, R., Muralidharan, V., Aron, A.R., 2020. Temporal cascade of frontal, motor and muscle processes underlying human action-stopping. eLife 9, e50371. URL: https://doi.org/10.7554/eLife.50371, doi:10.7554/eLife.50371. publisher: eLife Sciences Publications, Ltd.

Kemp, S.A., Bazin, P.L., Miletić, S., Boag, R.J., Keuken, M.C., Hinder, M.R., Forstmann, B.U., 2024. The ageing stopping network: Regional and network changes in the IFG, preSMA, and STN across the adult lifespan. URL: https://www.biorxiv.org/content/10.1101/2024.08.13.607702v1, doi:10.1101/2024.08.13.607702. pages: 2024.08.13.607702 Section: New Results.

Kenemans, J.L., 2015. Specific proactive and generic reactive inhibition. Neuroscience & Biobehavioral Reviews 56, 115–126. URL: http://www.sciencedirect.com/science/article/pii/S0149763415001657, doi:10.1016/j.neubiorev.2015.06.011.

Kibleur, A., Gras-Combe, G., Benis, D., Bastin, J., Bougerol, T., Chabardès, S., Polosan, M., David, O., 2016. Modulation of motor inhibition by subthalamic stimulation in obsessive-compulsive disorder. Translational Psychiatry 6, e922–e922. URL: https://www.nature.com/articles/tp2016192, doi:10.1038/tp.2016.192. number: 10 Publisher: Nature Pub-lishing Group.

Kim, Y., Choi, J., Kim, B., Park, Y., Cha, J., Choi, J., Han, S., 2024. Investigating the relationship between CSAT scores and prefrontal fNIRS signals during cognitive tasks using a quantum annealing algorithm. Scientific Reports 14, 19760. URL: https://www.nature.com/articles/s41598-024-70394-7, doi:10.1038/s41598-024-70394-7. publisher: Nature Publishing Group.

Ko, Y.T., Cheng, S.K., Juan, C.H., 2015. Voluntarily-generated unimanual preparation is associated with stopping success: evidence from LRP and lateralized mu ERD before the stop signal. Psychological Research 79, 249–258. doi:10.1007/s00426-014-0567-3.

Ko, Y.T., Miller, J., 2013. Signal-related contributions to stopping-interference effects in selective response inhibition. Experimental Brain Research 228, 205–212. URL: https://doi.org/10.1007/s00221-013-3552-y, doi:10.1007/s00221-013-3552-y.

Kohl, S., Hannah, R., Rocchi, L., Nord, C.L., Rothwell, J., Voon, V., 2019. Cortical paired associative stimulation influences response inhibition: Cortico-cortical and cortico-subcortical networks. Biological Psychiatry 85, 355–363. URL: http://www.sciencedirect.com/science/article/pii/S0006322318314057, doi:10.1016/j.biopsych.2018.03.009.

Lee, H.W., Lu, M.S., Chen, C.Y., Muggleton, N.G., Hsu, T.Y., Juan, C.H., 2016. Roles of the pre-SMA and rIFG in conditional stopping revealed by transcranial magnetic stimulation. Behavioural Brain Research 296, 459–467. URL: http://www.sciencedirect.com/science/article/pii/S016643281530156X, doi:10.1016/j.bbr.2015.08.024.

Li, C.S.R., Huang, C., Constable, R.T., Sinha, R., 2006. Imaging response inhibition in a stop-signal task: neural correlates independent of signal monitoring and post-response processing. The Journal of Neuroscience: The Official Journal of the Society for Neuroscience 26, 186–192. doi:10.1523/JNEUROSCI.3741-05.2006.

van der Linden, W.J., 2006. A lognormal model for response times on test items. Journal of Educational and Behavioral Statistics 31, 181–204. URL: https://doi.org/10.3102/10769986031002181, doi:10.3102/10769986031002181.

Lo, S., Andrews, S., 2015. To transform or not to transform: Using generalized linear mixed models to analyse reaction time data. Frontiers in Psychology 6. URL: https://www.frontiersin.org/articles/10.3389/fpsyg.2015.01171/full, doi:10.3389/fpsyg.2015.01171.

Logan, G.D., 1981. Attention, automaticity, and the ability to stop a speeded choice response, in: Long, J.D., Baddeley, A. (Eds.), Attention and performance IX. Erlbaum, Hillsdale, NJ, pp. 205–222.

Logan, G.D., Cowan, W.B., 1984. On the ability to inhibit thought and action: A theory of an act of control. Psychological Review 91, 295–327. doi:10.1037/0033-295X.91.3.295.

Love, J., Dropmann, D., Selker, R., 2022. The jamovi project. URL: https://www.jamovi.org.

MacDonald, H.J., McMorland, A.J.C., Stinear, C.M., Coxon, J.P., Byblow, W.D., 2017. An activation threshold model for response inhibition. PLoS ONE 12, e0169320. URL: https://www.ncbi.nlm.nih.gov/pmc/articles/PMC5235378/, doi:10.1371/journal.pone.0169320.

MathWorks, 2017. MATLAB and Statistics Toolbox R2017b.

Matzke, D., Verbruggen, F., Logan, G.D., 2018. The Stop-Signal Paradigm, in: Stevens’ Handbook of Experimental Psychology and Cognitive Neuroscience. John Wiley & Sons, Ltd, pp. 1–45. URL: https://onlinelibrary.wiley.com/doi/abs/10.1002/9781119170174.epcn510.

Messel, M.S., Raud, L., Hoff, P.K., Skaftnes, C.S., Huster, R.J., 2019. Strategy switches in proactive inhibitory control and their association with task-general and stopping-specific networks. Neuropsychologia 135, 107220. URL: http://www.sciencedirect.com/science/article/pii/S0028393218303671, doi:10.1016/j.neuropsychologia.2019.107220.

Meyer, H., Bucci, D., 2016. Neural and behavioral mechanisms of proactive and reactive inhibition. Learning and Memory 23, 504–514. doi:10.1101/lm.040501.115.

Miller, J., 1991. Reaction time analysis with outlier exclusion: bias varies with sample size. The Quarterly Journal of Experimental Psychology. A, Human Experimental Psychology 43, 907–912. doi:10.1080/14640749108400962.

Morís Fernández, L., Vadillo, M.A., 2020. Flexibility in reaction time analysis: many roads to a false positive? Royal Society Open Science 7, 190831. URL: https://www.ncbi.nlm.nih.gov/pmc/articles/PMC7062108/, doi:10.1098/rsos.190831.

Nambu, A., Tokuno, H., Takada, M., 2002. Functional significance of the cortico–subthalamo–pallidal ‘hyperdirect’ pathway. Neuroscience Research 43, 111–117. URL: https://www.sciencedirect.com/science/article/pii/S0168010202000275, doi:10.1016/S0168-0102(02)00027-5.

Nieuwenhuis, S., Forstmann, B.U., Wagenmakers, E.J., 2011. Erroneous analyses of interactions in neuroscience: A problem of significance. Nature Neuroscience 14, 1105–1107. URL: http://www.nature.com/articles/nn.2886, doi:10.1038/nn.2886.

Obeso, I., Robles, N., Muñoz-Marrón, E., Redolar-Ripoll, D., 2013. Dissociating the role of the pre-SMA in response inhibition and switching: a combined online and offline TMS approach. Frontiers in Human Neuroscience 7. URL: https://www.frontiersin.org/journals/human-neuroscience/articles/10.3389/fnhum.2013.00150/full, doi:10.3389/fnhum.2013.00150. publisher: Frontiers.

Obeso, I., Wilkinson, L., Teo, J.T., Talelli, P., Rothwell, J.C., Jahanshahi, M., 2017. Theta burst magnetic stimulation over the pre-supplementary motor area improves motor inhibition. Brain Stimulation 10, 944–951. URL: http://www.sciencedirect.com/science/article/pii/S1935861X17307994, doi:10.1016/j.brs.2017.05.008.

Pani, P., Giarrocco, F., Bardella, G., Brunamonti, E., Ferraina, S., 2022. Reply to: Hannah et al. (2021) Commentary: ‘Does action-stopping involve separate pause and cancel processes? A view from premotor cortex’: Action-stopping models must consider the role of the dorsal premotor cortex. Cortex 152, 160–163. URL: https://www.sciencedirect.com/science/article/pii/S001094522200106X, doi:10.1016/j.cortex.2022.03.017.

Peirce, J., Gray, J.R., Simpson, S., MacAskill, M., Höchenberger, R., Sogo, H., Kastman, E., Lindeløv, J.K., 2019. PsychoPy2: Experiments in behavior made easy. Behavior Research Methods 51, 195–203. URL: https://doi.org/10.3758/s13428-018-01193-y, doi:10.3758/s13428-018-01193-y.

Pinti, P., Tachtsidis, I., Hamilton, A., Hirsch, J., Aichelburg, C., Gilbert, S., Burgess, P.W., 2020. The present and future use of functional near-infrared spectroscopy (fNIRS) for cognitive neuroscience. Annals of the New York Academy of Sciences 1464, 5–29. URL: https://www.ncbi.nlm.nih.gov/pmc/articles/PMC6367070/, doi:10.1111/nyas.13948.

Puri, R., Nikitenko, T., Kemp, S.A., 2018. Using transcranial magnetic stimulation to investigate the neural mechanisms of inhibitory control. Journal of Neurophysiology 120, 1587–1590. URL: https://www.physiology.org/doi/10.1152/jn.00366.2018, doi:10.1152/jn.00366.2018.

Rae, C.L., Hughes, L.E., Weaver, C., Anderson, M.C., Rowe, J.B., 2014. Selection and stopping in voluntary action: A meta-analysis and combined fMRI study. NeuroImage 86, 381–391. URL: http://www.sciencedirect.com/science/article/pii/S1053811913010240, doi:10.1016/j.neuroimage.2013.10.012.

Raez, M., Hussain, M., Mohd-Yasin, F., 2006. Techniques of EMG signal analysis: detection, processing, classification and applications. Biological Procedures Online 8, 11–35. URL: https://www.ncbi.nlm.nih.gov/pmc/articles/PMC1455479/, doi:10.1251/bpo115.

Rahman, M.A., Siddik, A.B., Ghosh, T.K., Khanam, F., Ahmad, M., 2020. A narrative review on clinical applications of fNIRS. Journal of Digital Imaging 33, 1167–1184. URL: https://www.ncbi.nlm.nih.gov/pmc/articles/PMC7573058/, doi:10.1007/s10278-020-00387-1.

Raud, L., Huster, R., Ivry, R., Labruna, L., Messel, M., Greenhouse, I., 2020. A single mechanism for global and selective response inhibition under the influence of motor preparation. Journal of Neuroscience 40, 7921–7935. doi:10.1523/JNEUROSCI.0607-20.2020.

Raud, L., Thunberg, C., Huster, R.J., 2022. Partial response electromyography as a marker of action stopping. eLife 11, e70332. URL: https://elifesciences.org/articles/70332, doi:10.7554/eLife.70332.

Rocha, G.S., Freire, M.A.M., Britto, A.M., Paiva, K.M., Oliveira, R.F., Fonseca, I.A.T., Araújo, D.P., Oliveira, L.C., Guzen, F.P., Morais, P.L.A.G., Cavalcanti, J.R.L.P., 2023. Basal ganglia for beginners: the basic concepts you need to know and their role in movement control. Frontiers in Systems Neuroscience 17, 1242929. URL: https://www.ncbi.nlm.nih.gov/pmc/articles/PMC10435282/, doi:10.3389/fnsys.2023.1242929.

Rolls, E.T., Joliot, M., Tzourio-Mazoyer, N., 2015. Implementation of a new parcellation of the orbitofrontal cortex in the automated anatomical labeling atlas. NeuroImage 122, 1–5. URL: https://www.sciencedirect.com/science/article/pii/S1053811915006953, doi:10.1016/j.neuroimage.2015.07.075.

Salomoni, S.E., Gronau, Q.F., Heathcote, A., Matzke, D., Hinder, M.R., 2023. Proactive cues facilitate faster action reprogramming, but not stopping, in a response-selective stop signal task. Scientific Reports 13, 19564. URL: https://www.nature.com/articles/s41598-023-46592-0, doi:10.1038/s41598-023-46592-0.

Salomoni, S.E., Weber, S., Hinder, M.R., 2024. An exploration of complex action stopping across multiple datasets: Insights into the mechanisms of action cancellation and re-programming. URL: https://www.biorxiv.org/content/10.1101/2024.11.03.621753v1, doi:10.1101/2024.11.03.621753. pages: 2024.11.03.621753 Section: New Results.

Schmidt, R., Berke, J.D., 2017. A Pause-then-Cancel model of stopping: evidence from basal ganglia neurophysiology. Philosophical Transactions of the Royal Society of London. Series B, Biological Sciences 372, 20160202. doi:10.1098/rstb.2016.0202.

Schroll, H., Hamker, F.H., 2013. Computational models of basal-ganglia pathway functions: focus on functional neuroanatomy. Frontiers in Systems Neuroscience 7. URL: https://www.frontiersin.org/journals/systems-neuroscience/articles/10.3389/fnsys.2013.00122/full, doi:10.3389/fnsys.2013.00122.

Schwarz, G., 1978. Estimating the dimension of a model. The Annals of Statistics 6, 461–464. URL: https://projecteuclid.org/journals/annals-of-statistics/volume-6/issue-2/Estimating-the-Dimension-of-a-Model/10.1214/aos/1176344136.full, doi:10.1214/aos/1176344136.

Sebastian, A., Forstmann, B.U., Matzke, D., 2018. Towards a model-based cognitive neuroscience of stopping – a neuroimaging perspective. Neuroscience & Biobehavioral Reviews 90, 130–136. URL: http://www.sciencedirect.com/science/article/pii/S014976341830040X, doi:10.1016/j.neubiorev.2018.04.011.

Sebastian, A., Konken, A.M., Schaum, M., Lieb, K., Tüscher, O., Jung, P., 2020. Surprise: unexpected action execution and unexpected inhibition recruit the same fronto-basal-ganglia network. The Journal of Neuroscience, JN–RM–1681–20URL: http://www.jneurosci.org/lookup/doi/10.1523/JNEUROSCI.1681-20.2020, doi:10.1523/JNEUROSCI.1681-20.2020.

Sebastian, A., Pohl, M.F., Klöppel, S., Feige, B., Lange, T., Stahl, C., Voss, A., Klauer, K.C., Lieb, K., Tüscher, O., 2013. Disentangling common and specific neural subprocesses of response inhibition. NeuroImage 64, 601–615. URL: http://www.sciencedirect.com/science/article/pii/S1053811912009305, doi:10.1016/j.neuroimage.2012.09.020.

Senkowski, D., Ziegler, T., Singh, M., Heinz, A., He, J., Silk, T., Lorenz, R.C., 2023. Assessing inhibitory control deficits in adult ADHD: A systematic review and meta-analysis of the stop-signal task. Neuropsy-chology Review URL: https://doi.org/10.1007/s11065-023-09592-5, doi:10.1007/s11065-023-09592-5.

Sharp, D.J., Bonnelle, V., De Boissezon, X., Beckmann, C.F., James, S.G., Patel, M.C., Mehta, M.A., 2010.Distinct frontal systems for response inhibition, attentional capture, and error processing. Proceedings of the National Academy of Sciences of the United States of America 107, 6106–6111. doi:10.1073/pnas.1000175107.

St George, R.J., Hinder, M.R., Puri, R., Walker, E., Callisaya, M.L., 2021. Functional near-infrared spectroscopy reveals the compensatory potential of pre-frontal cortical activity for standing balance in young and older adults. Neuroscience 452, 208–218. doi:10.1016/j.neuroscience.2020.10.027.

St George, R.J., Jayakody, O., Healey, R., Breslin, M., Hinder, M.R., Callisaya, M.L., 2022. Cognitive inhibition tasks interfere with dual-task walking and increase prefrontal cortical activity more than working memory tasks in young and older adults. Gait & Posture 95, 186–191. URL: https://www.sciencedirect.com/science/article/pii/S0966636222001217, doi:10.1016/j.gaitpost.2022.04.021.

Sánchez-Carmona, A.J., Albert, J., Hinojosa, J.A., 2016. Neural and behavioral correlates of selective stopping: Evidence for a different strategy adoption. NeuroImage 139, 279–293. URL: http://www.sciencedirect.com/science/article/pii/S1053811916302920, doi:10.1016/j.neuroimage.2016.06.043.

Tatz, J.R., Soh, C., Wessel, J.R., 2021. Common and unique inhibitory control signatures of action-stopping and attentional capture suggest that actions are stopped in two stages. Journal of Neuroscience 41, 8826–8838. doi:10.1523/JNEUROSCI.1105-21.2021.

Team, R.C., 2021. R: A language and environment for statistical computing. URL: https://cran.r-project.org.

Thunberg, C., Wiker, T., Bundt, C., Huster, R.J., 2024. On the (un)reliability of common behavioral and electrophysiological measures from the stop signal task: Measures of inhibition lack stability over time. Cortex 175, 81–105. URL: https://www.sciencedirect.com/science/article/pii/S0010945224000479, doi:10.1016/j.cortex.2024.02.008.

Tzourio-Mazoyer, N., Landeau, B., Papathanassiou, D., Crivello, F., Etard, O., Delcroix, N., Mazoyer, B., Joliot, M., 2002. Automated anatomical labeling of activations in SPM using a macroscopic anatomical parcellation of the MNI MRI single-subject brain. NeuroImage 15, 273–289. URL: https://www.sciencedirect.com/science/article/pii/S1053811901909784, doi:10.1006/nimg.2001.0978.

Ulrich, R., Miller, J., 1994. Effects of truncation on reaction time analysis. Journal of Experimental Psychology. General 123, 34–80. doi:10.1037//0096-3445.123.1.34.

Verbruggen, F., Aron, A.R., Band, G.P., Beste, C., Bissett, P.G., Brockett, A.T., Brown, J.W., Chamberlain, S.R., Chambers, C.D., Colonius, H., Colzato, L.S., Corneil, B.D., Coxon, J.P., Dupuis, A., Eagle, D.M., Garavan, H., Greenhouse, I., Heathcote, A., Huster, R.J., Jahfari, S., Kenemans, J.L., Leunissen, I., Li, C.S.R., Logan, G.D., Matzke, D., Morein-Zamir, S., Murthy, A., Paré, M., Poldrack, R.A., Ridderinkhof, K.R., Robbins, T.W., Roesch, M., Rubia, K., Schachar, R.J., Schall, J.D., Stock, A.K., Swann, N.C., Thakkar, K.N., van der Molen, M.W., Vermeylen, L., Vink, M., Wessel, J.R., Whelan, R., Zandbelt, B.B., Boehler, C.N., 2019. A consensus guide to capturing the ability to inhibit actions and impulsive behaviors in the stop-signal task. eLife 8, e46323. URL: https://elifesciences.org/articles/46323, doi:10.7554/eLife.46323.

Verbruggen, F., Chambers, C.D., Logan, G.D., 2013. Fictitious inhibitory differences: how skewness and slowing distort the estimation of stopping latencies. Psychological Science 24, 352–362. doi:10.1177/0956797612457390.

Vink, M., Kaldewaij, R., Zandbelt, B.B., Pas, P., du Plessis, S., 2015. The role of stop-signal probability and expectation in proactive inhibition. The European Journal of Neuroscience 41, 1086–1094. doi:10.1111/ejn.12879.

Wadsley, C.G., Cirillo, J., Nieuwenhuys, A., Byblow, W.D., 2022. Stopping interference in response inhibition: Behavioral and neural signatures of selective stopping. The Journal of Neuroscience 42, 156–165. URL: x, doi:10.1523/JNEUROSCI.0668-21.2021.

Wadsley, C.G., Cirillo, J., Nieuwenhuys, A., Byblow, W.D., 2023. A global pause generates nonselective response inhibition during selective stopping. Cerebral Cortex, bhad239 URL: https://academic.oup.com/cercor/advance-article/doi/10.1093/cercor/bhad239/7216778, doi:10.1093/cercor/bhad239.

Wadsley, C.G., Greenhouse, I., 2024. Failures to launch preclude response inhibition. Trends in Cognitive Sciences URL: https://www.sciencedirect.com/science/article/pii/S1364661324000548, doi:10.1016/j.tics.2024.03.001.

Watanabe, H., Shitara, Y., Aoki, Y., Inoue, T., Tsuchida, S., Takahashi, N., Taga, G., 2017. Hemoglobin phase of oxygenation and deoxygenation in early brain development measured using fNIRS. Proceedings of the National Academy of Sciences 114, E1737–E1744. URL: https://www.pnas.org/doi/full/10.1073/pnas.1616866114, doi:10.1073/pnas.1616866114.

Watanabe, T., Hanajima, R., Shirota, Y., Tsutsumi, R., Shimizu, T., Hayashi, T., Terao, Y., Ugawa, Y., Katsura, M., Kunimatsu, A., Ohtomo, K., Hirose, S., Miyashita, Y., Konishi, S., 2015. Effects of rTMS of pre-supplementary motor area on fronto basal ganglia network activity during stop-signal task. Journal of Neuroscience 35, 4813–4823. URL: https://www.ncbi.nlm.nih.gov/pmc/articles/PMC6705371/, doi:10.1523/JNEUROSCI.3761-14.2015.

Weber, S., Salomoni, S.E., Hinder, M.R., 2024a. Selective cancellation of reactive or anticipated movements: Differences in speed of action reprogramming, but not stopping. Cortex 177, 235–252. URL: https://www.sciencedirect.com/science/article/pii/S0010945224001485, doi:10.1016/j.cortex.2024.05.010.

Weber, S., Salomoni, S.E., Kilpatrick, C., Hinder, M.R., 2023. Dissociating attentional capture from action cancellation during the inhibition of bimanual movement. Psychophysiology, e14372 URL: https://onlinelibrary.wiley.com/doi/10.1111/psyp.14372, doi:10.1111/psyp.14372.

Weber, S., Salomoni, S.E., St George, R.J., Hinder, M.R., 2024b. Stopping speed in response to auditory and visual stop signals depends on go signal modality. Journal of Cognitive Neuroscience 36, 1395–1411. doi:10.1162/jocn_a_02171.

van den Wildenberg, W.P.M., Ridderinkhof, K.R., Wylie, S.A., 2022. Towards conceptual clarification of proactive inhibitory control: A review. Brain Sciences 12, 1638. URL: https://www.ncbi.nlm.nih.gov/pmc/articles/PMC9776056/, doi:10.3390/brainsci12121638.

Zhu, T., Cao, C., Coxon, J., Sack, A.T., Leunissen, I., 2024. Using TMS-EEG to study the intricate interplay between GABAergic inhibition and glutamatergic excitation during reactive and proactive motor inhibition. URL: https://www.biorxiv.org/content/10.1101/2024.09.05.610957v1, doi:10.1101/2024.09.05.610957. pages: 2024.09.05.610957 Section: New Results.

Zimeo Morais, G.A., Balardin, J.B., Sato, J.R., 2018. fNIRS Op-todes’ Location Decider (fOLD): a toolbox for probe arrangement guided by brain regions-of-interest. Scientific Reports 8, 3341. URL: https://www.ncbi.nlm.nih.gov/pmc/articles/PMC5820343/, doi:10.1038/s41598-018-21716-z.

